# A foundation language model to decipher diverse regulation of RNAs

**DOI:** 10.1101/2024.10.12.617732

**Authors:** Hanwen Zhou, Yue Hu, Yulong Zheng, Jiefu Li, Jielong Peng, Jiang Hu, Yun Yang, Guoqing Zhang, Zefeng Wang

**Affiliations:** Bio-Med Big Data Center, Chinese Academy of Sciences Key Laboratory of Computational Biology, Shanghai Institute of Nutrition and Health, University of Chinese Academy of Sciences, Chinese Academy of Sciences, Shanghai 200031, China; School of Life Science, Guangming Advanced Research Institute, Southern University of Science and Technology, Shenzhen, Guangdong, China; School of Medicine, Department of Pharmacy, University of North Carolina at Chapel Hill, Chapel Hill, NC, USA; CirCode Biomedicine Inc., Shanghai, China

**Keywords:** RNA regulation, language model, splicing, translation, degradation

## Abstract

RNA metabolism is tightly regulated by *cis*-elements and *trans*-acting factors. Most information guiding such regulation is encoded in RNA sequences. Considering the similarities in semantic and syntactic features between RNAs and human language, we developed LAMAR, a transformer-based foundation <u>la</u>nguage <u>m</u>odel for RN<u>A</u> <u>r</u>egulation, to decipher general rules underlying RNA processing. The model was pretrained on approximately 15 million sequences from both genome and transcriptome of 225 mammals and 1569 viruses, and further fine-tuned with labeled datasets for various tasks. The resulting fine-tuned models outperformed the state-of-the-art methods in predicting mRNA translation efficiency and mRNA half-life, while achieving comparable accuracy to specifically designed methods in predicting splice sites of pre-mRNAs and internal ribosome entry sites. Our results indicated that a single foundation language model is applicable in the comprehensive analysis of different aspects of RNA regulation, providing new insight into the design and optimization of RNA drugs.

## Introduction

As an important component of the central dogma, RNAs play various roles in multiple biological processes, ranging from the templates of protein synthesis ^1^ to the regulators of gene expression ^2^. The cellular processes of RNA are regulated by its *cis*-regulatory elements and *trans*-acting factor ^3, 4^, the information of which are encoded in RNA sequences spanning evolutionary diversity. In addition, the regulation of RNAs is often affected by RNA structures, which can either directly determine the RNA metabolism or affect the accessibility of RNA *cis*-elements recognized by the cognate RNA binding protein (RBPs) ^5, 6^. Deciphering the regulatory rules of RNA will provide new insights into molecular mechanisms and further direct the designs of new RNA therapeutics.

With the advancement of biochemical and bioinformatic technologies, researchers have previously developed different computational tools to analyze the regulatory maps of RNAs ^7–11^. While these well-designed algorithms were specifically developed for different aspects of RNA regulation, a unified framework to learn the diverse regulation of RNA is lacking. The large language models (LLM) have demonstrated that unsupervised pretraining of a vast corpuses of text can significantly advance our comprehension ^12^ and generative capabilities of language ^13^. The pretrained model serves as a foundation platform that can be further fine-tuned to solve multiple downstream tasks. In light of the similarity of semantic and syntactic features between RNA sequences and human language, some studies explored the potential of language models to capture the essence of RNA. For example, RNA-MSA ^14^ and RNAErnie ^15^ were developed to predict the secondary structures of non-coding RNAs. RNA-FM was pretrained on non-coding RNAs and fine-tuned to solve multiple tasks including prediction of RNA tertiary structures and binding sites of RNA binding proteins ^16^. However, current models were mostly pretrained on non-coding RNA sequences, and thus the extracted sequence features are mainly related to the functions of non-coding RNAs. As a result, these models may be biased to the highly structured non-coding RNAs, therefore may not be suitable for downstream tasks that interpret different aspects of RNA regulation. Meanwhile, the gene expression undergoes multi-layer regulation at the RNA level, including alternative splicing, polyadenylation, localization, modification, mRNA translation and degradation. The regulatory rules are presumably encoded in the sequence, theoretically dictating the structure and the RBP binding profile. However, uncovering such hidden information is technically challenging due to the complexity, diversity and context-dependency nature of RNA regulation. Additionally, the evolution information can provide crucial insights, helping to distinguish conserved regulatory elements and to reveal their functional significance.

To address these challenges, we developed a foundation <u>la</u>nguage <u>m</u>odel for RN<u>A</u> <u>r</u>egulation (LAMAR) by unsupervised pretraining using approximately 15 million RNA sequences with multiple functions across mammalians. LAMAR can effectively distinguish between functional regions or transcripts with distinct functions within the representation space, indicating that it has successfully captured the functional attributes solely from sequences. By leveraging LAMAR as a foundation platform, we were able to fine-tune this model with labeled datasets across a range of tasks including supervised modeling of splice sites, RNA stability, mRNA translation efficiency, and internal ribosome entry sites (IRESs) for non-canonical translation. Our results showed that the performances of the fine-tuned LAMARs were better than or comparable with the state-of-the-art methods that were specially designed for each task, which demonstrated the power of LLM in dissecting the diverse functions of RNAs using only the primary sequences and provided new clues for underlying cellular mechanisms.

## Results

### The pretrained RNA foundation model

We developed an RNA foundation model, LAMAR, by first pretraining with large-scale unlabeled RNA sequences and subsequently fine-tuning with task-specific datasets to solve multiple downstream tasks across different layers of RNA regulation. To efficiently learn the sequence features and evolutionary information of RNA, the ∼21 million sequences from all types of genes and transcripts across 225 mammals from 15 suborders (Supplementary Figure 1A) and 1569 viruses were downloaded and processed as training corpus from RefSeq and RNACentral databases (Figure 1A, also see methods). We further reduced the data redundancy by selecting representative sequences from different clusters (Supplementary Figure 1B, also see methods), resulting in a final dataset of 15 million sequences that encompass a broad range of species and length (Figure 1B). These sequences included the mRNAs, non-coding RNAs and annotated genes from all these species, covering around 267 billion nucleotide tokens for the training of LAMAR. In the pretraining stage, we used the mask learning strategy in which a portion of the nucleotides were masked and then predicted by contextual sequences, enabling the model to learn the interactions between nucleotides (Figure 1C). Considering the sequence length of genes/transcripts and the available computational resources, we pretrained two models with the contextual length of up to 2048 and 4096 tokens, named LAMAR-2k and LAMAR-4k, resulting in two encoder models that are linked to different task-specific prediction heads (Figure 1C). With more than 50% accuracy in reconstructing masked tokens, the pretrained models successfully learned the exact contextual information after 250 thousand steps (Supplementary Figure 2). The task-specific predictor was then stacked on the encoder for each downstream tasks within the fine-tuning. To examine the performance and generality of LAMAR, we applied this RNA foundation model in different downstream tasks ranging from the RNA splicing, translation and degradation (Figure 1D).

**Figure 1.**
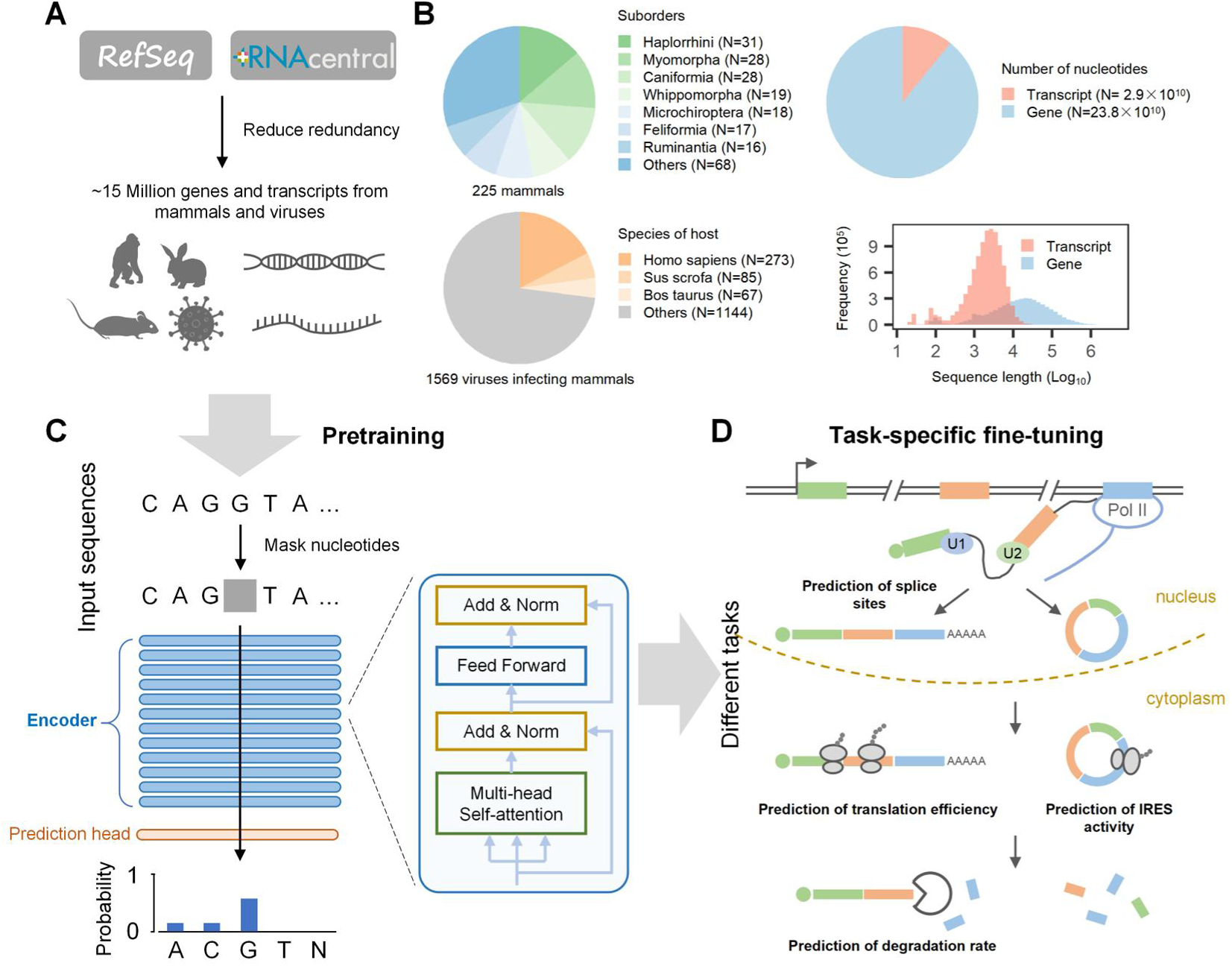
Pretraining an RNA foundation model for multilayer regulation of RNA. **(A)** Schematic for processing of the pretraining dataset. **(B)** The statistics of pretraining dataset. Left, the taxonomic makeup of the mammals (top) and the viruses (bottom) used for pretraining. Right, the number of the nucleotide (top) and sequence length (bottom) of the genes and transcripts in the pretraining dataset. **(C)** In the pretraining stage, LAMAR learns the sequence features of RNA by predicting the likelihoods of the masked tokens in the sequence. LAMAR comprises of 12 transformer layers and a task-specific prediction head. **(D)** LAMAR is fine-tuned for the downstream tasks including supervised prediction of splice site, translation efficiency, degradation rate and internal ribosome entry site (IRES).

### Extraction of RNA intrinsic features with pretrained LAMAR

To investigate whether the model can learn the intrinsic biochemical properties of RNA during the pretraining stage, we first compared the hidden embeddings of the four nucleotides between the untrained and pretrained model (Figure 2A). In this test, each nucleotide was represented as a single vector by averaging the output embeddings of nucleotides in 500 randomly selected human transcripts (including both mRNAs and long non-coding RNAs). We then used the principal component analysis to project the representative vectors into two-dimensional space. We found that, while the nucleotides from the untrained model are scattered randomly, the four different nucleotides were clustered together after the pretraining. Additionally, we found that, in the first principal component (PC1), the adenine and guanine (purines) are closer to each other, while the cytosine and thymine (pyrimidines) are closely clustered. This result indicated that the model had somehow learned the biochemical properties of different nucleotides from large-scale unlabeled sequences (Figure 2A). The distances of hidden embeddings were presented in Supplementary Figure 3.

**Figure 2.**
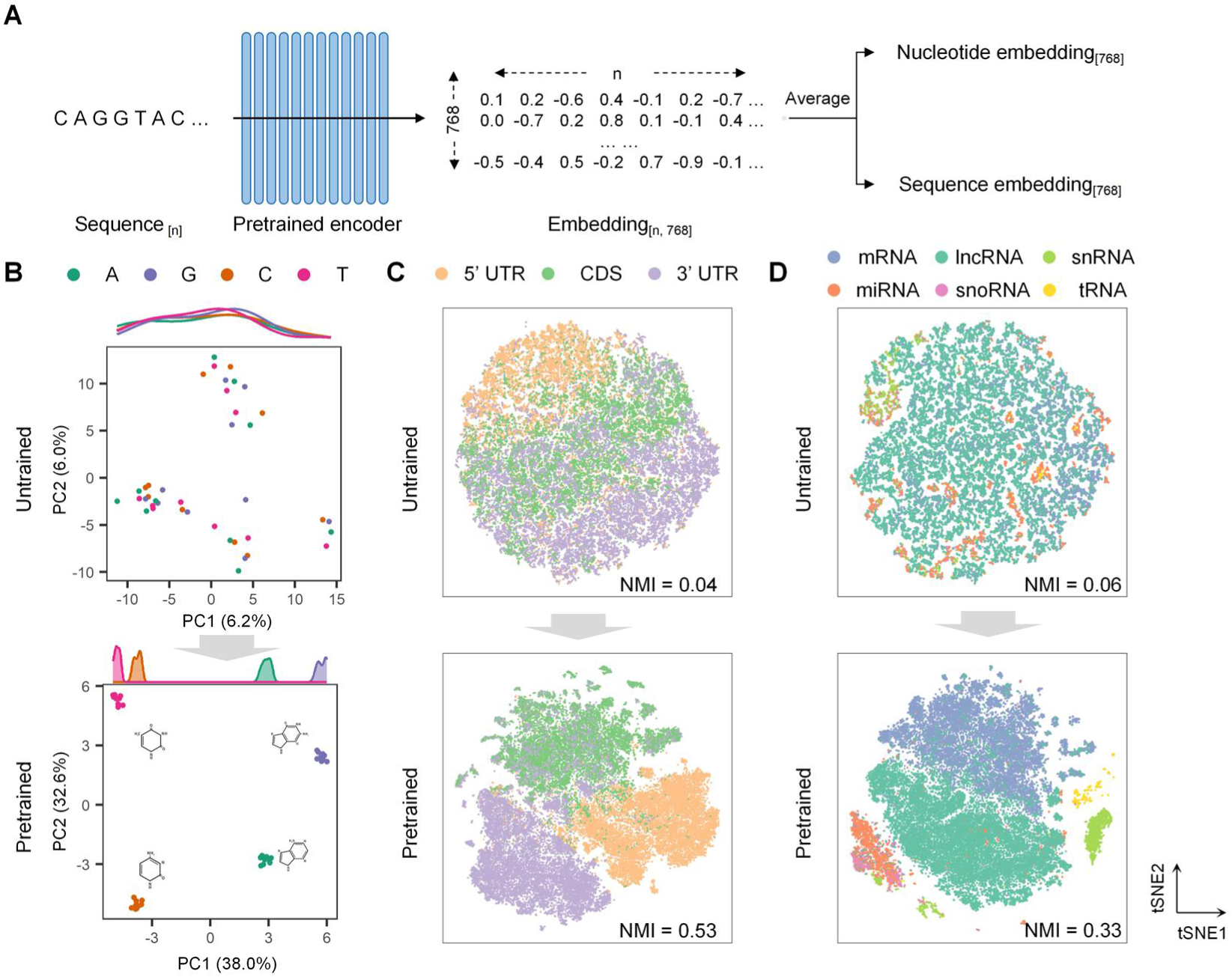
LAMAR learned the biological properties of nucleotides and the function features of RNAs. **(A)** Schematic for the representative embedding by LAMAR. The LAMAR pretrained encoder of represents the RNA sequence of *n* nucleotides as the hidden embedding with the shape of *n* × 768. The hidden embedding was further averaged as the representative embedding of the nucleotide and RNA for visualization of their feature space. **(B-D)** Visualization of the embeddings from the pretrained model (top) and the untrained model (bottom). NMI, normalized mutual information. **(B)** The PCA plot visualizing the embeddings of four classes of nucleotides. **(C)** The tSNE plot visualizing the embeddings of functional regions (5′ UTR, CDS and 3′ UTR). **(D)** The tSNE plot visualizing the embeddings of RNAs (mRNA, lncRNA, snRNA, miRNA, snoRNA and tRNA).

Encouraged by the ability of LAMAR in discerning different RNA bases, we further examined if the pretraining of LAMAR could, to a degree, extract some functional features of RNAs. We first test if the pretraining of LAMAR can distinguish different functional regions within mRNA (CDS *vs* UTRs) by representing different mRNA fragments as vectors using the untrained and the pretrained models, and subsequently projecting them in a two-dimensional plot using t-Distributed Stochastic Neighbor Embedding (tSNE) (Figure 2B). We found that the RNA sequences with same functions were more likely to be clustered together after pretraining (Normalized Mutual Information, NMI=0.53) compared to the random scattering of the same sequences in untrained model (NMI=0.04) (Figure 2B). The similar results were also obtained for different functional types of RNAs (NMI=0.060 in untrained model *vs* NMI=0.327 after pretraining) (Figure 2C). These results indicate that the model has successfully extracted the intrinsic functional features of RNA in self-supervised pretraining with unlabeled sequences.

### The fine-tuned LAMAR accurately predicts RNA splice sites

The majority of mammalian genes are transcribed as pre-mRNAs that contain intervening introns and exons. During the process of pre-mRNA splicing, the spliceosomes precisely recognize pairs of splice sites (SS) and subsequently remove the introns, and the adjacent exons are joined together to produce mature mRNAs ^17^. In addition to authentic SS, pre-mRNAs contain numerous “decoy” SS that have similar consensus motifs but are really spliced, probably because introns contain large number of splicing silencer sequences to suppress the use of decoy SS ^18^. Therefore distinguishing the real SS among the large number of “decoy” sites become a central question in splicing regulation ^19^, and the accurate prediction of SS from human transcripts is also crucial for identifying disease-causing genetic variants ^20^. Various computational tools have been previously developed to predictively identify splice sites within pre-mRNA sequences, among which the best performance was achieved by a CNN-based method SpliceAI ^8^.

To examine if the LAMAR can be used to distinguish the authentic SS among the decoy sites, we clustered the real SS and the decoy sites (non-splicing GT and AG sites) by projecting their representative vectors into the two-dimensional space using tSNE. We found that while the 5′ splice sites and other GT dinucleotides could not be classified clearly for the untrained model (NMI=0.00), the pretrained LAMAR can clearly separate these two types of sites with good clustering efficiency (NMI=0.49) (Figure 3B). The same result was also observed for 3′ splice sites (NMI=0.00 in untrained model *vs* NMI=0.45 in pretrained LAMAR), indicating that the pretrained model has successfully learned the contexts of splice sites, which may be used in the subsequent fine-tuning.

**Figure 3.**
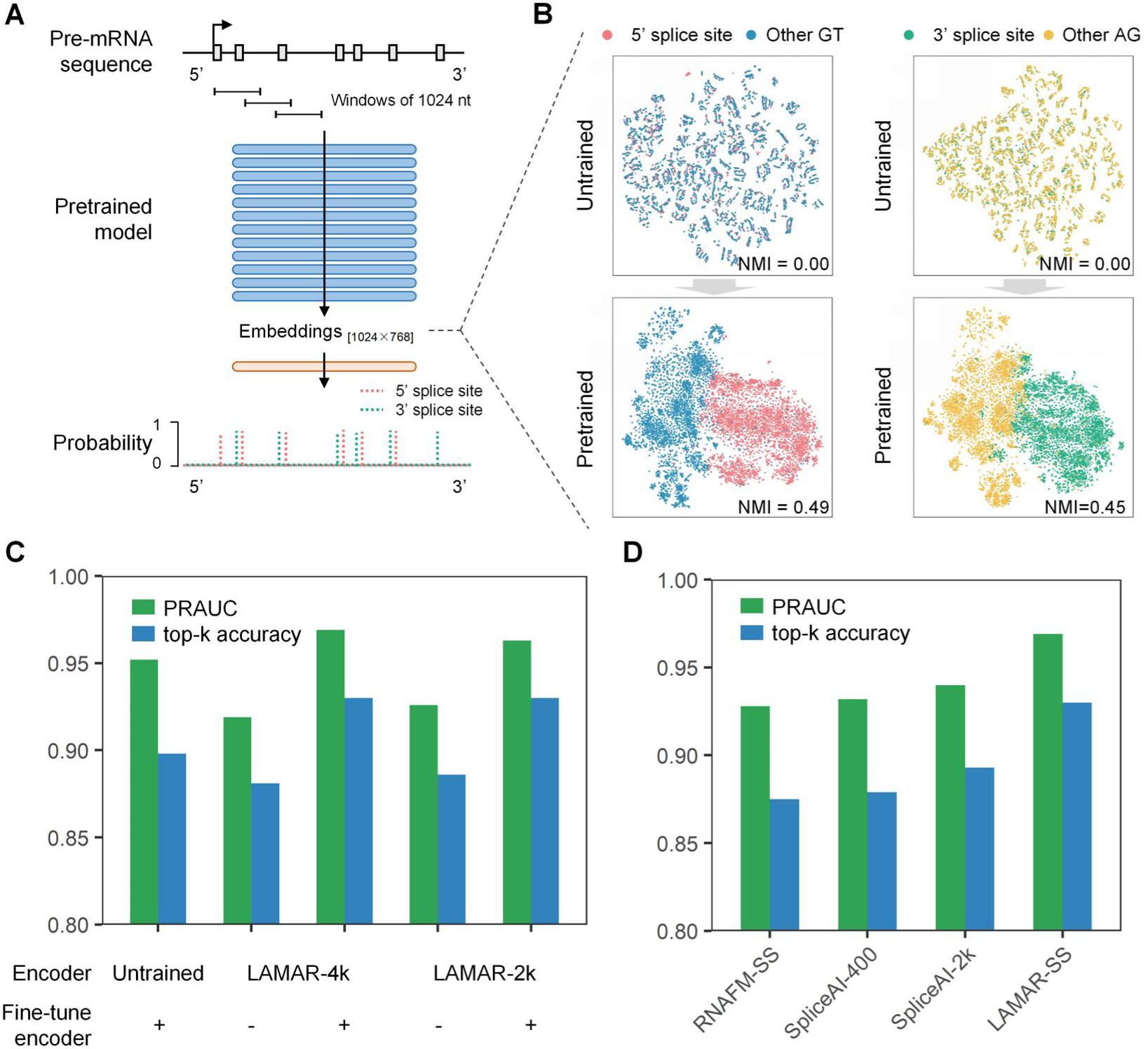
Prediction of splice sites in pre-mRNAs. **(A)** Schematic for prediction of splice sites in pre-mRNAs by fine-tuned LAMAR. First, LAMAR represents the pre-mRNA segments of 1024 nt as the hidden embeddings of the shape 1024 × 768. Then, the hidden embeddings were input into the prediction head to predict the probability that a nucleotide is a 5′ splice site, a 3′ splice site or neither. **(B)** The tSNE plots visualizing the hidden embeddings of the splice sites and non-splice sites output from the untrained model (top) and the pretrained model (bottom). **(C)** The top-k accuracy and PRAUC of LAMAR in predicting splice sites, evaluated under different fine-tuning strategies. **(D)** The top-k accuracy and PRAUC of LAMAR-SS and baseline methods (including SpliceAI sub-models and RNA-FM-SS) in predicting splice sites.

We further fine-tuned LAMAR to predict the 5′ and 3′ splice sites for each transcript in human pre-mRNA sequences. We used the same datasets consistent with SpliceAI and set the contextual length as 1024 tokens for the task-specific fine-tuning (Figure 3A). Two fine-tuning strategies were tested for both LAMAR-2k and LAMAR-4k: 1) fine-tuning frozen LAMAR (i.e., only the parameters in prediction head are updated, and the LAMAR encoder is not changed during the fine-tuning stage), and 2) fine-tuning all the parameters of LAMAR (i.e., both the prediction head and LAMAR encoder are updated during the fine-tuning, see methods). As a control, we also fine-tuned the untrained LAMAR with the same prediction head for comparison. We used precision-recall area under curve (PRAUC) and top-k accuracy as model metrics, and found that the second strategy using global optimization outperformed the control (untrained LAMAR) and the first strategy (frozen LAMAR), indicating the effectiveness of pretraining and the parameter optimization of the pretrained model for prediction (Figure 3C). It is worth noting that the frozen pretrained model with only a simple prediction head (i.e., the first strategy) could also achieve pretty good result (PRAUC=0.926 and top-k accuracy=0.886 for LAMAR-2k), suggesting that the pretrained LAMAR has already captured many sequence features of the splice sites. We defined the best prediction model, the LAMAR-4k with all parameters fine-tuned, as LAMAR-SS and compared it with baseline methods including SpliceAI and RNA-FM. To control the effect of contextual length on model performance, the sub-models spliceAI-400 and spliceAI-2k were selected which predicted the splicing sites based on flanking 200nt and 1000nt sequences.

The results show that LAMAR-SS achieved better results than the other baseline methods in splice site predictions as judged by both PRAUC and top-k accuracy (Figure 3D). In particular, LAMAR-SS outperformed RNA-FM-SS by 0.041, spliceAI-400 by 0.037 and spliceAI-2k by 0.029 in terms of PRAUC (outperformed RNA-FM-SS by 0.055, spliceAI-400 by 0.051 and spliceAI-2k by 0.037 in terms of top-k accuracy), showing the powerful performance of pretrained model.

### Fine-tune LAMAR to decipher the translation regulation by 5′ UTR

The translation efficiency of mRNA is critical for the development of mRNA drugs, which is tightly regulated by general and specific translation factors inside cells ^21^. To gain insight on how the RNA sequence may affect translation efficiency, we next examined if the LAMAR model can be applied to predict the regulation of translation. It is well accepted that the 5′ untranslated region (5′ UTR) of mRNA plays a major role in the regulation of mRNA translation ^22^. The structures and *cis*-elements located in sequences including hairpins and Kozak sequences control the translation efficiency (TE) of mRNA. In addition, recent experiments with massively parallel reporter assays have suggested that the translation regulatory elements are very prevalent throughout human 5′ UTRs ^23, 24^.

Using the available experimental data from high-throughput profiling of human 5′ UTR, we fine-tuned LAMAR to predict the translation efficiency of human mRNAs based on 5′ UTR sequences (Figure 4A). The 10,903 transcripts from human embryonic kidney (HEK) 293T cell line were selected for training ^25^. To select the best fine-tuned strategy, we compared the same fine-tuned strategies in prediction of splice sites. The results showed that fine-tuned all the parameters of LAMAR-2k achieved the best result (mean squared error MSE=0.409 and Spearman correlation coefficient=0.652), which we named it as LAMAR-TE (Supplementary Figure 4). Since UTR-LM outperformed than classical methods in prediction of translation efficiency of 5′ UTRs ^26^, we selected UTR-LM and RNA-FM as benchmarks. The results of 10-fold cross validation showed that LAMAR-TE outperformed RNA-FM and UTR-LM by 7% and 18% for the Spearman correlation coefficient (Figure 4B), and by 9% and 14% for the MSE respectively (Figure 4C). UTR-LM was pretrained on sequences and secondary structures of 5′ UTRs using approximately 1 million parameters. Therefore the superior performance of LAMAR-TE compared to UTR-LM may suggest that our model probably learned the structural information of 5′ UTRs exclusively from sequences with much more parameters.

**Figure 4.**
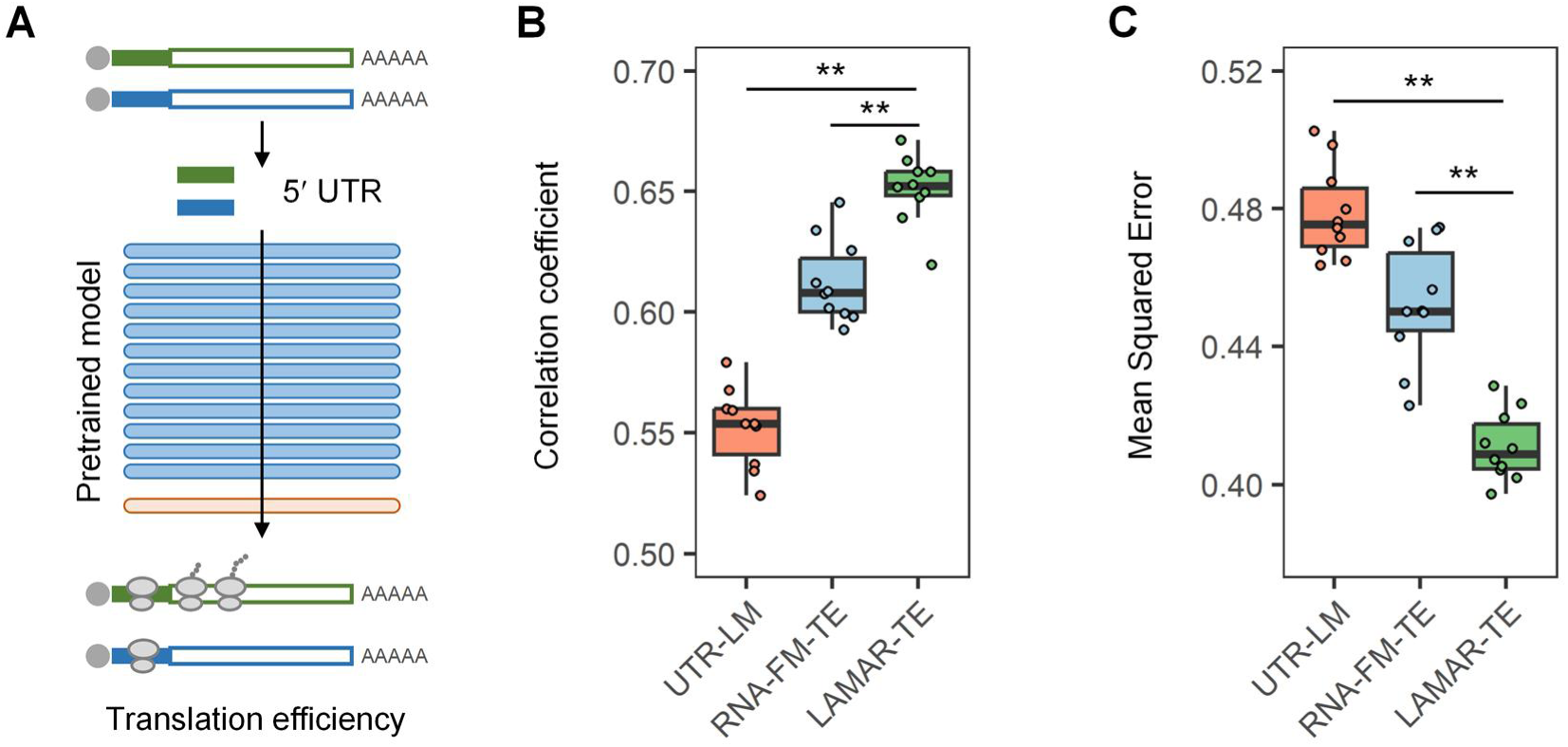
Prediction of translation efficiencies of mRNAs by 5′ UTRs. **(A)** Schematic for prediction of translation efficiencies of mRNAs based on 5′ UTRs by fine-tuned LAMAR. **(B, C)** The Spearman correlation coefficient and mean squared error between observed and predicted translation efficiency for LAMAR-TE and baseline methods, including RNA-FM-TE and UTR-LM. The *P* values were calculated with Wilcoxon signed rank test. **, P value < 0.01.

### Predict the effect of 3′ UTRs on RNA degradation using fine-tuned LAMAR

In general, mRNAs are degraded at the end of their live cycles by exo/endonucleases and associated factors ^27^. The sequences in 3′ untranslated regions (3′ UTRs) are known to play major roles in regulating various degradation pathways by recruiting miRNAs or RBPs ^28^. To study the effects of the *cis*-regulatory elements and genetic variants of 3′ UTRs on mRNA stability, the degradation rates (DR) of native and mutated 3′ UTR sequences have been measured in several massive parallel reporter assays ^29, 30^.

In this section, we fine-tuned our LAMAR to predict the half-lives of mRNAs containing different 3′ UTRs (Figure 5A). We used the data from a massive parallel reporter assay that measures the effect of 1,967 human 3′ UTRs on the half-lives of eGFP mRNAs containing different 3′ UTRs ^30^. The model performance was tested by predicting the effect of 3′ UTRs on mRNA half-lives with ten-fold cross-validation. We compared the performances of various fine-tuning strategies, and found that the full-parameter fine-tuned LAMAR-4k, named LAMAR-DR, have achieved the best performance in predicting half-lives of the reporter mRNAs (MSE=0.176 and Spearman correlation coefficient=0.647) (Supplementary Figure 5A, B). The fine-tuned frozen LAMAR-4k achieved the modest result (MSE=0.223 and Spearman correlation coefficient=0.477), showing that LAMAR-4k extracted the sequence features of 3′ UTRs from unlabeled pretraining. As a comparison with baseline method, we fine-tuned the RNA-FM using the same dataset and strategy, which we named as RNA-FM-DR. We found that, while both foundation models showed ability in predicting mRNA half-lives, LAMAR-DR outperformed RNA-FM-DR by 8.0% for correlation coefficient (Figure 5B) and by 7.4% for MSE (Figure 5C), suggesting that LAMAR has a superior performance in model generalization.

**Figure 5.**
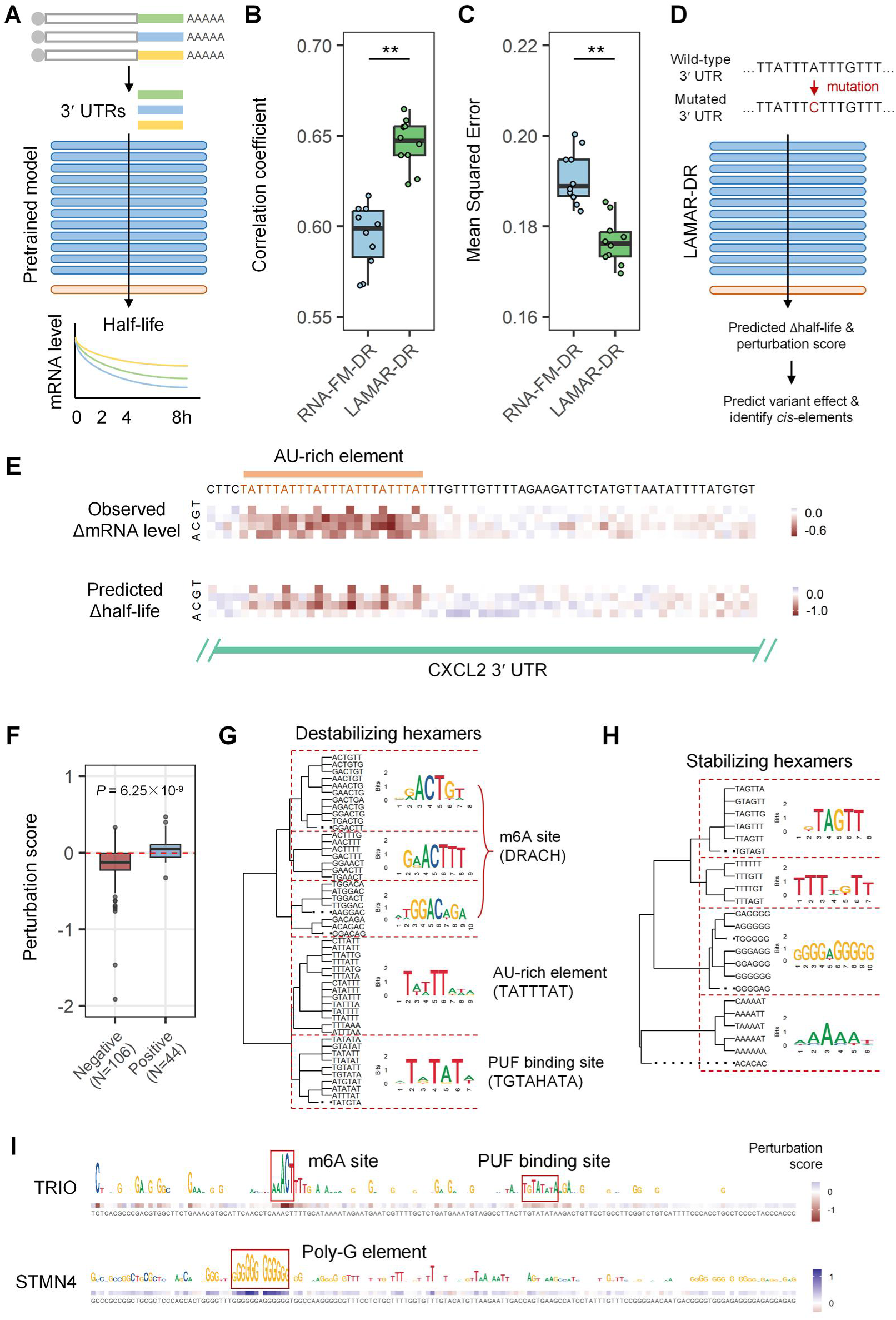
Prediction of degradation rates of mRNAs by 3′ UTRs. **(A)** Schematic for fine-tuning LAMAR to predict the half-lives of mRNAs with different 3′ UTRs. **(B, C)** The Spearman correlation coefficient and mean squared error between observed and predicted half-lives for LAMAR-DR and RNA-FM-DR. **(D)** Each nucleotide of 3′ UTR was mutated *in silico* and the half-lives of mutated sequences were predicted by LAMAR-DR. The Δhalf-lives between the mutated and wild-type 3′ UTRs were computed to predict the mutation effect, and the perturbation scores were analyzed to identify *cis*-regulatory elements. **(E)** The observed (top) and predicted (bottom) mutation effects of 3′ UTR segment of CXCL2. The nucleotides of the AU-rich element were colored in orange. **(F)** The distribution of perturbation scores of the short elements which positively and negatively regulate the half-life of mRNA. **(G, H)** The identified destabilizing and stabilizing hexamers which were clustered into 5 and 4 classes by the polygenetic tree, respectively. The names of reported motifs were listed at the right of the clusters. **(I)** The examples of the LAMAR-identified destabilizing and stabilizing motifs were presented by the profile of perturbation score in 3′ UTR segments of TRIO and STMN4 genes. *P* values were calculated with Wilcoxon signed rank test. **, *P* value < 0.01.

We next evaluated LAMAR-DR by examining whether the model predicts how the 3′ UTR mutations may affect mRNA stability (Figure 5D). We used the 3′ UTR segment of CXCL2 mRNA that had been systematically mutated in a near saturated fashion ^30^. For each mutation, we predicted the half-lives of the mutant mRNA and the natural sequence using LAMAR-DR, and defined their difference as the variant effect. We compared the predicted variant effect with the experimental results measuring the effect of each CXCL2 mutation on the mRNA level ^30^, and found that the LAMAR-DR prediction was largely correlated with the experimental measurement of mRNA levels (Figure 5E). In addition, LAMAR-DR successfully identified a strong degradation element, AU-rich element, in the upstream of 3′ UTR (Figure 5E). For all the mutations on CXCL2 3′ UTR, the Pearson correlation coefficient between predicted variant effect and experimental results was 0.58 (Supplementary Figure 5C). Considering the difference in experimental measurement between mRNA half-lives and steady state levels ^30^, the correlation between LAMAR-DR prediction and experimental measurement is reasonably strong.

To examine the sequence features used by LAMAR-DR in predicting RNA degradation, we applied perturbation analysis on each nucleotide to measure how they affect the prediction of RNA stability (Figure 5D). We define the perturbation score of a sequence as the differences of the LAMAR-DR score between the wildtype sequences and the average scores of their mutations (see methods), and such *in silico* mutation analysis can generate the regulatory elements in 3′ UTR that either promote or inhibit degradation of mRNAs. Previously, a massively parallel functional assay was conducted using the >450 kilobases of 3′ UTR sequences from >2000 human genes to measure their effect on mRNA stability, identifying 150 short elements that positively or negatively regulate mRNA stability ^30^. We used the LAMAR-DR to computed the average perturbation scores of these 150 regulatory elements ^30^, and found that the positive regulators of mRNA stability indeed showed a significantly higher score than the negative regulators (Figure 5F). In addition, the LAMAR-DR predicted regulatory effects are positively correlated with observed values (Spearman correlation coefficient=0.58, *P*=5.5 × 10^-15^, Supplementary Figure 5D), demonstrating the effectiveness and potential of our method in discovering new regulatory elements.

To predict new *cis*-elements that may regulate mRNA degradation, we computed the LAMAR-DR perturbation scores of all hexamers in human 3′ UTRs, and analyzed the hexamers with the highest and lowest scores (top or bottom 1%) as the stabilizing and destabilizing elements that can be clustered into several groups to obtain the consensus motifs (Figure 5G). As the examples, two 3′ UTR containing the stabilizing or destabilizing sites were included together with their perturbation scores (Figure 5I). Interestingly, we found that the destabilizing motifs contain several known motifs to promote mRNA degradation, such as the AU-rich elements (UAUUUAU), PUF binding elements (TGTAHATA) and m6A sites (DRACH) (Figure 5G), confirming the reliability of our prediction. However, most of the stabilizing motifs, include GA-rich, U-rich and A-rich elements, seem to be novel (Figure 5H), although the AAAAU motif resembles the known binding site of IGF2BP3 that is involved in regulating mRNA stability ^4, 31^.

### The fine-tuned LAMAR for IRES prediction

The internal ribosome entry sites (IRESs) can drive cap-independent translation of eukaryotic mRNAs, which is essential for the viral mRNAs lacking 5′ caps or the translation of certain mRNAs under stressful conditions ^32^. Canonical IRESs were mostly identified from the viral genomes, and some IRESs were reported in the endogenous human genes ^32^. Previously several algorithms, such as IRESPred ^33^, IRESfinder ^34^ and IRESpy ^35^, have been built to computationally predict IRESs. However, their performances are not satisfactory and unable to produce coherent and reliable results for IRES prediction, probably due to data noise or insufficient training data. Recently, a large number of IRESs were identified from high-throughput screening of human and viral sequences ^36–38^, most of which have not been annotated and analyzed thoroughly to generate a predictive model. On the other hand, identification of IRESs is important for learning the mechanisms of cap-independent translation and improving circRNA-based biomedical engineering.

With an RNA foundation model, we next seek to fine-tune LAMAR to predict new sequences with IRES activity (Figure 6A). We collected 1,901 experimentally validated IRESs from the three databases (IRESite ^39^, IRESbase ^40^ and RFAM ^41^) as the positive training sequences (Figure 6B), and used the structural RNAs randomly sampled from RFAM database as the negative control sequences (see methods). We achieved the best result by fine-tuning all the parameters of LAMAR-2k (named as LAMAR-IRES), which was slightly better than LAMAR-4k (Supplementary Figure 6). Since the previous IRES prediction methods were trained on short sequences that are inherently noisy, we did not use them as benchmark, and instead fine-tuned RNA-FM as the benchmark. We found that both of fine-tuned models from RNA-FM and LAMAR achieved the good AUC around 0.98 (Figure 6C), suggesting that the RNA foundational model have a strong power in extracting the features of known IRES datasets to make a great prediction.

**Figure 6.**
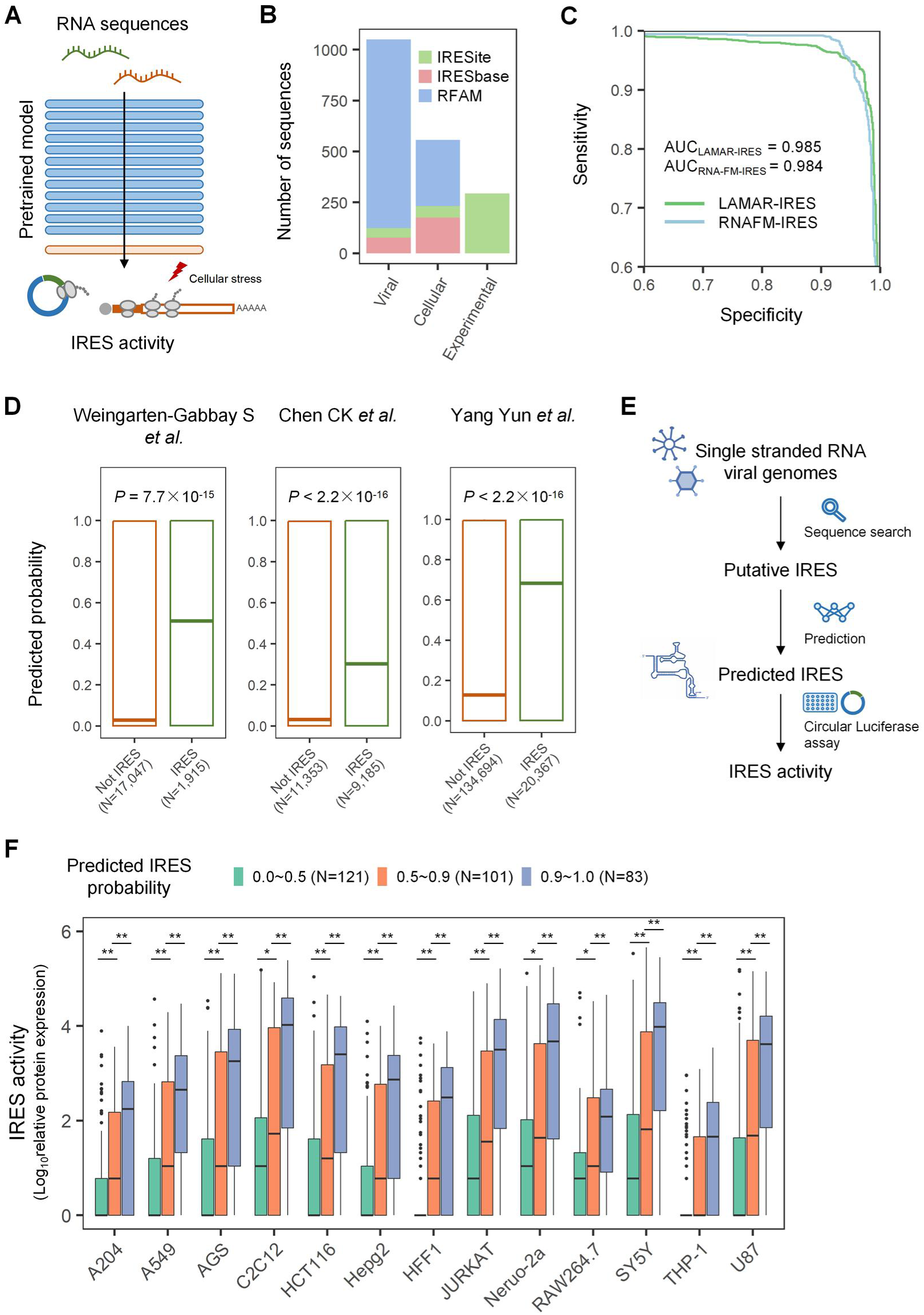
Prediction of IRES. **(A)** Schematic for fine-tuning LAMAR to predict IRESs. **(B)** The IRESs collected from IRESite, IRESbase and RFAM databases were used as training data. **(C)** The ROC curves of LAMAR-IRES and RNA-FM-IRES. **(D)** The predicted probabilities of oligos in three high-throughput experimental libraries screening IRES. **(E)** Schematic of *in silico* screening of IRES. The putative IRESs were extracted from positive-sense single-strand RNA viruses and their probabilities were predicted by LAMAR-IRES. The IRES activities of the candidates were subsequently validated by the circRNA-based luciferase assays. **(F)** The IRES activities of candidate sequences classified by LAMAR-IRES scores were tested in 13 cell lines. *P* values were calculated with Wilcoxon rank sum test. *, *P* value < 0.05; **, *P* value < 0.01.

To examine the performance of LAMAR-IRES in predicting the novel IRESs obtained from independent experimental screen, we used LAMAR-IRES to classify the recently identified IRESs from the high throughput screens of a library derived from viral and cellular mRNAs ^36–38^. We found that the predicted probabilities of putative IRESs are significantly higher than those of remaining sequences, revealing the *in silico* screening ability of LAMAR-IRES (Figure 6D).

The circular RNAs (circRNAs) have been found to be able to translated into functional protein in human cells ^42, 43^, and thus can served as a new generation of mRNA therapy due to its high stability and low immunogenicity ^44–46^. Since circRNAs lack free 5′ or 3′ ends, they can only be translated through a cap-independent fashion driven by IRESs or IRES-like elements ^43, 47^. Therefore, the identification of highly efficient IRESs is critical for the development of circRNA-based therapy, as the IRES-mediated translation is generally less efficient compared to the canonical cap-dependent translation. To address this issue, we next applied the LAMAR-IRES to screen 5′ UTRs of viral genomes in order to predict new IRES sequences that are highly active in promoting circRNA translation (Figure 6E). Specifically, we applied a series of filters to selected 305 annotated 5′ UTRs or upstream regions from the genomes of positive-sense single-stranded RNA viruses, and used LAMAR-IRES to score the probability of the candidate sequences (see methods). We classified the candidates into three groups based on the predicted probabilities: non-IRESs (probability<0.5), low-activity IRESs (0.5≤probability<0.9) and high-activity IRESs (0.9≤probability). To validate our prediction, each group of sequences were subsequently inserted into the *in vitro* synthesized circRNAs, and their protein expression in 13 frequently-used cell lines were measured by quantitative luciferase assays (see methods, Supplementary Table 2). The activity of IRES was assessed by calculating the relative expression levels of circRNA and modified mRNA, both containing the identical reporter gene. We compared the activities between three IRES groups and found that the high-activity IRESs exhibited significantly higher activities than low-activity IRESs, which in turn were significantly higher than those of non-IRES in each cell line (Figure 6F), suggesting that our model integrated the causal relationship between RNA sequence and IRES activity. In addition, we observed mostly consistent results for these sequences in 13 cultured cell lines, suggesting that the prediction of LAMAR-IRES can be validated in different cell contexts. Furthermore, we examined 14 highly active IRESs (probability>0.9) predicted by LAMAR-IRES, and found that the protein expression level driven by these IRESs surpassed the modified mRNA in specific cell lines. For instance, the circRNA containing the IRES from parabovirus A4 (probability=0.91) exhibited 2.44-fold higher protein expression than the mRNA with same ORF sequences in C2C12 cells. However, four IRESs with high activities were predicted with low probability (probability<0.5), suggesting that the sensitivity of LAMAR-IRES prediction should be further improved with better training data quality.

## Discussion

To study the multi-layer regulation of RNA, we build a foundation language model, LAMAR, which can extract semantic features of RNAs by unsupervised pretraining on unlabeled transcripts with diverse functions across mammals and viruses. The representation space learned from LAMAR reflected the biological functions of RNA in multiple levels including nucleotide, regulatory region and entire sequence. We further fine-tuned the pretrained LAMAR with labeled datasets for several downstream tasks, including prediction of splice sites, mRNA translation efficiency, mRNA degradation rate and IRES activity, and demonstrated that the foundation model showed high versatility and accuracy on handing these tasks. In particular, LAMAR achieved better or comparable performance than the baseline methods in nearly all tasks evaluated, reflecting the remarkable generality of LAMAR in modeling RNA regulation. The future application of LAMAR may be expanded to study other processes of RNA regulation including alternative polyadenylation, transportation/localization, editing and modification. More importantly, LAMAR can be applied to predict mutational effects of *cis*-regulatory elements and to identify novel regulatory elements, demonstrating the potential of this model in revealing new biological mechanisms and biomedical engineering.

LAMAR was pretrained on a diverse dataset comprising genomic and transcriptomic sequences from a wide range of species, ensuring its adaptability for downstream tasks. Since the RNA regulation includes multiple layers, the generality of LAMAR makes it easier to achieve robust performance across diverse regulatory mechanisms. For instance, gene sequences play a critical role in understanding RNA splicing regulation, as the selection of splice sites is influenced by regulatory elements within both introns and exons ^17^. The pretraining with both genomic and transcriptomic sequences could give additional attention to the regions of mature transcripts that play more important biological roles. During the pretraining process, we clustered the genome and transcriptome sequences to reduce redundancy. However, certain groups of RNAs may lack sufficient representation after clustering. Therefore, fine-tuning the model is necessary to improve predictions for these underrepresented groups.

LAMAR shared a similar architecture with some of the benchmark models like RNA-FM and RNA-Ernie, however was pretrained on different corpus using datasets containing a huge number of pre-mRNAs, mature mRNAs, and non-coding RNAs. The diverse types of training data in LAMAR may be responsible for its superior performance in downstream tasks related to co-/post-transcriptional regulation compared to the other methods that were mainly pretrained on the non-coding RNAs with high content of secondary structure. It is generally accepted that RNA structures play critical roles in the gene expression. However, compared to the number of available sequences, only a small number of accurate RNA structures are currently available, which are insufficient to pretrain a large model. In addition, RNA structures are very dynamic and affected by their binding factors as well as the cellular conditions ^48^, and thus direct pretraining with RNA sequences and their structures would mislead the learning of complex features. Even in the cases where the secondary structures of RNAs can be predicted, the accuracy of structure prediction is not equally reliable for different types of RNAs ^49^, introducing uncontrolled bias during the pretraining with a large set of diverse RNAs. For all these reasons, we decide to not specially consider the RNA structure during the pretraining stage, and instead acknowledge that the structural information has already been embedded in the sequences and thus may be sensed/learned by a powerful model. It may worth trying to combine the sequence embeddings and structural information for the task-specific fine-tuning in the future applications.

Since LAMAR uses the architecture of transformer encoders containing many self-attention layers, the number of parameters is proportional to the square of the input length. Due to the limitation of graphic memory, some ultra-long RNA sequences have to be split into the overlapping short segments (with the length of 2k or 4k) during pretraining and fine-tuning, which may lead to the loss of distant coevolutionary information in RNA. Although the regulations of RNAs is mostly determined by the interactions within local context, there are some functional attributes of RNAs, such as translation efficiency and degradation rate, are dependent on entire sequences including coding and untranslated regions ^50^. Therefore, training with partial sequence elements may reduce the model performance because the long-range interactions are likely be diminished during the pretraining. Recently several frameworks have been designed for ultra-long sequences ^51^, which will be explored in the updated version of LAMAR.

Although LAMAR has ∼85 million parameters, the pretraining loss curve shows that it may underfit the pretraining dataset (Supplementary Figure 2). The evolvement of large language models in deciphering human languages ^52^ and protein structure ^53, 54^ suggests that the model performance is roughly proportional to the model size, implying the future potential of LAMAR with an expanding model scale and data diversity. The future expansion of such model probably requires the exploration of more efficient architectures and/or new computational algorithm and hardware.

Developing a foundation model to understand RNA regulation and design functional RNA components is an important application of artificial intelligence in the field of RNA biology and therapeutics. We believe this study provide the opportunity of a general-purpose model which can integrate fundamental properties of RNA and design novel regulatory elements with desired functions. The future versions will be optimized toward more efficient architectures, robust reasoning and enhanced generation, and we will also include different combinations of training dataset for additional pretraining.

## Methods

### 1. Model architecture

In this work, we developed a foundation language model, LAMAR, to study diverse regulation of RNAs. LAMAR takes the RNA sequence as input and represents the nucleotides and RNAs. In LAMAR, the RNA sequence *x* of *x* nucleotides (nt) is tokenized as tokens [*x*_1_, *x*_2_, …, *x*_*x*_], where *x*_*i*_ ∈ V = {‘A’, ‘T’, ‘C’, ‘G’, ‘N’}. The special tokens 〈*cls*〉 and 〈*eos*〉 are added to the beginning and end of the tokens, which denotes the start and end of the sequence. Each token *x*_*i*_ is further embedded as *e*_*i*_ ∈ ℝ^*h*^*D*, and the sequence *x* is represented as the embedding matrix *e* = [*e*_*cls*_, *e*_1_, *e*_2_, …, *e*_*x*_, *e*_*eos*_] ∈ ℝ(*x*2)^×ℎ^*D*, where hidden size ℎ_*D*_ is set to be 768. Subsequently, the embedding *e* is further input into a multi-layer bidirectional Transformer encoder similar with ESM-2 ^54^ to get the final hidden embedding matrix ℎ = [ℎ_*cls*_, ℎ_1_, ℎ_2_, …, ℎ_*x*_, ℎ_*eos*_] ∈ ℝ(*x*_2_)^×ℎ^*D*. At last, the final hidden embedding ℎ representing the entire sequence is fed into the task-specific prediction head for different downstream tasks.

Specifically, the multi-layer bidirectional Transformer encoder contains 12 layers, each of which comprises a self-attention layer with 12 heads and a feed forward layer with intermediate size of 3072. To enhance the model performance on long sequences, the rotary position embedding is applied to integrate positional information into the learning process of the model ^55^.

### 2. Pretraining dataset

The pretrained datasets of LAMAR derived from four sources: 1) Protein coding and non-coding transcripts of mammals from RefSeq ^56^; 2) Non-coding transcripts of mammals from RNACentral ^57^; 3) Reference genome of mammals from NCBI Genome; 4) Reference genome of viruses that infect mammals from NCBI Genome.

RefSeq (Release 221) contained 13,319,105 transcripts of 223 mammalian species in 15 suborders. The sequences were downloaded as fna.gz format from website https://ftp.ncbi.nlm.nih.gov/refseq/release/vertebrate_mammalian/. RNACentral (Release 23) contained 2,649,775 non-coding transcripts from 214 mammalian species. The sequences could be downloaded from https://ftp.ebi.ac.uk/pub/databases/RNAcentral/releases/23.0/sequences/. As of Dec 14 2023, NCBI Genome contained 222 mammalian reference genomes annotated by RefSeq. The sequences and annotations were respectively downloaded as fna.gz and gff.gz from the links on the metadata file (https://ftp.ncbi.nlm.nih.gov/genomes/refseq/vertebrate_mammalian/assembly_summary.txt). The 6,033,179 reference genes were extracted from the reference genomes by “bedtools” ^58^ with the genome annotations. As of Jan 4 2024, NCBI Virus contained 15,037 viral reference genomes, 1579 of which infected mammals were included into pretrain datasets (https://ftp.ncbi.nlm.nih.gov/genomes/refseq/viral/assembly_summary.txt). The 19,007 reference genes were extracted by “bedtools” ^58^ with the genome annotations. The above data were combined into the initial corpus, comprising 15,455,049 transcripts and 5,832,792 genes in total.

For the initial corpus, the sequences of the length fewer than 15 nt were filtered. The IUPAC ambiguity codes in sequences were replaced by N. To reduce the data redundancy, we used MMseq2 ^59^ to respectively cluster transcripts and genes based on the sequence identity. Specifically, Clustering was performed with a minimum threshold of 80% sequence identity and 80% coverage, and the longest sequences are used as the representative sequence in each cluster (cluster mode=2, identity=0.8, coverage=0.8). As a result, we obtained 6,673,915 transcript and 4,557,965 gene clusters. During pretraining, 99% of transcript and gene clusters were randomly selected to pretrain the model, while the remaining clusters were reserved for evaluation. The representative sequences and 50% of the member sequences from each training cluster were included into pretraining corpus, with an upper limit of 200 members per cluster. In addition, the validation corpus was composed of the representative sequences of testing cluster. In all, the training corpus encompassed a total of 15,000,406 sequences, which accounted for approximately 267 billion tokens. The testing corpus comprised 112,797 sequences, providing representative samples for validation.

### 3. Pretraining LAMAR

In the pretraining stage, we trained the model with masked language modeling ^12, 54^. We first randomly masked 15% of tokens in each sequence. For each masked token, we replaced it with the 〈*mask*〉 token 80% of the time, the random token 10% of the time and the unchanged token 10% of the time. After masking, we input the masked sequence into the model and used the final hidden embedding ℎ to predict the labels of the masked tokens. The categorical cross-entropy loss between predicted probabilities and the actual labels of the masked tokens was used to pretrain the model ^12^. The loss function was defined as:

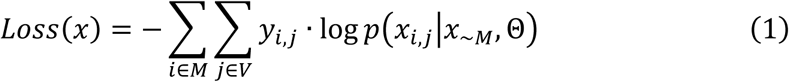

where *M* and ∼*M* represent the positions of masked nucleotides and unmasked nucleotides, respectively. Θ represents the trainable parameters of the model and *px*_*i*,_*x*_∼*M*_, Θ represents the probability of predicting the *i*_*j*ℎ_ masked token as class *j* based on all the unmasked tokens. *y*_*i*,_ represents whether *i*_*j*ℎ_ masked token belongs to class *j*, if so, *y*_*i*,_=1, otherwise, *y*_*i*,_=0. We pretrained two models with contextual lengths of 2048 and 4096 tokens respectively, to accommodate different cases, named as LAMAR-2k and LAMAR-4k. When the length of the pretraining sequences exceeded these limits, they were segmented into fragments shorter than the specified contextual length. The overlap of 256 and 512 tokens between adjacent fragments were ensured to maintain contextual information. For the evaluation phase, we randomly extracted fragments with contextual length from testing sequences that exceeded the pretraining length. Any testing sequences with more than 10% Ns were excluded. To speed up evaluating the model, 10,000 sequences were randomly sampled from the testing corpus to compute the accuracy of predicting the labels of masked tokens.

To pretrain the models stably, we set the batch sizes as 512 and 256 sequences (around 1M tokens) respectively for LAMAR-2k and LAMAR-4k. For optimization of model pretraining, we utilized the AdamW optimizer ^60^ (β_1_ = 0.9, β_2_ = 0.98, ε = 10) and ε_2_ weight decay of 0.01. In the pretraining stage, the learning rate was warmed up during the first 10,000 steps to the peak value 1e-4 and then decayed in remaining steps. To speed up pretraining without sacrificing accuracy, the models were pretrained with FP16 mixed precision with Apex and FlashAttention library ^61^. Our models underwent 250,000 training steps to reach convergence, taking around 260 hours on 8 A800 80GB graphic process units.

### 4. RNA representation by LAMAR

We took the final hidden embedding of LAMAR to represent the nucleotide, splice site, functional region (UTR and CDS) and RNA for visualization of feature space.

For the representative embeddings of four nucleotides, we averaged the final embeddings of the same class of nucleotides in RNA sequences *X* as formula:

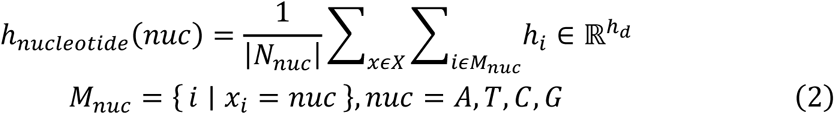

where *M*_*nuc*_ represents all the positions of the nucleotide *nuc* in sequence *x* and *N*_*nuc*_ represents the number of the positions of the nucleotide *nuc* in total sequences *X*. To compare the nucleotide embeddings of pretrained and untrained models, we sampled 500 human mRNAs and non-coding RNAs for 10 times to compute the representative embeddings of nucleotides. We only sampled one RNA from each gene and included sequences with lengths shorter than 4096 nt.

The splice site, functional region and RNA can be seen as sequence of different length. For each sequence, the final representative embedding is obtained by averaging the embeddings of all nucleotides in the sequence, as described by the formula (3).

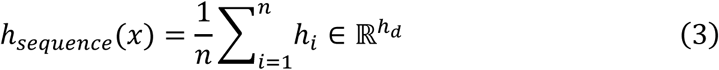

For the representative embeddings of splice sites and non-splice sites, we averaged the final embeddings of two bases of the specific sites in pre-mRNA sequences. We first recorded the positions of splice sites with RefSeq annotation file and used “bedtools” ^58^ to extract the pre-mRNA sequences from human genome. For each pre-mRNA, the non-splicing GT and AG sites with the same number of splice sites were sampled as non-splice sites. The 1,000 pre-mRNAs with lengths shorter than 14,500 nt were sampled and the 9,464 pairs of splice sites and non-splice sites were selected, respectively.

We averaged the final hidden embeddings of the total positions of different functional regions (UTR and CDS) in human mRNAs as their representative embeddings. The mRNAs annotated by GENCODE V43 containing 5¢ UTR, CDS and 3¢ UTR were first selected. We only sampled one mRNA from each gene to avoid the bias from high identities between mRNA alternative isoforms, and included 14,984 sequences with lengths shorter than 4096 nt. The hidden embeddings of functional regions were computed by untrained or pretrained LAMAR for comparison.

For the representative embeddings of transcripts with different RNA types, we averaged the final embeddings of all the positions in transcripts. All the transcripts annotated by GENCODE V43 were first selected. We only sampled one transcript from each gene and included transcripts with lengths shorter than 4096 nt. We selected the functional types with the number more than 100, including mRNA, lncRNA, snRNA, miRNA, snoRNA and tRNA. The representative vectors of transcripts were computed by untrained or pretrained LAMAR for comparison.

### 5. Fine-tuning LAMAR

For different downstream tasks, the fine-tuning can be categorized into token-level and sequence-level. For token-level tasks including prediction of splice site of pre-mRNA, the entire embedding ℎ from the encoder is fed into the prediction head which contains one linear layer. As for sequence-level tasks including prediction of translation efficiency, half-life and IRES, only the representation of the 〈*cls*〉 token ℎ_*cls*_ is input into the prediction head which contains two linear layers. Compared to the pretraining stage, fine-tuning is inexpensive, which takes several hours to finish training.

During fine-tuning, we explored two strategies. The first one is to fine-tune the frozen pretrained LAMAR, that is, the weights of the encoder are untrainable and only the task-specific head is fine-tuned. In this condition, the fine-tuning is speeded up and the memory usage is reduced. The second one is to fine-tune all the parameters end-to-end, which fits LAMAR to specific datasets, improving the model performance compared to the first strategy in all the tasks.

To evaluate the performance of the fine-tuned models for prediction of splice site and IRES, we used the specific strategy (see section 6.1 and 6.4). As for prediction of translation efficiency and degradation rate, we used cross-validation to evaluate the performance of the fine-tuned models. Specifically, 10% of the dataset was served as independent validation set and the remaining data was used for 10-fold cross-validation. All the hyperparameters of the fine-tuned models were the same as the pretrained model, except batch size, learning rate, warmup ratio and training steps. The warmup ratio was set to be 0.05. Since the model performance was influenced by different hyperparameters, the optimal hyperparameters (batch size, learning rate and training steps) are decided by grid searching (Supplementary table 1). To speed up the query of hyperparameters, we first fine-tuned the model with one random subset and ranked the combinations of hyperparameters by the performance. Further, we evaluated the performance of the top10 combinations of hyperparameters by 10-fold cross-validation. At last, we used the median performance of the 10 sub-models on the independent validation set as the final evaluation (Supplementary table 1). The model was fine-tuned with FP32 precision for better performance on the Sugon Z-100 16GB and Tesla V100 32GB clusters of graphic process units.

### 6. Downstream tasks

#### 6.1. Prediction of splice sites

We fine-tuned LAMAR with pre-mRNA sequences to predict the splice site. In the pre-mRNA sequence, each nucleotide is classified as the first site of 5¢ splice site (1), the second site of 3¢ splice site (2) or neither (0). Therefore, predicting the splice sites of human pre-mRNAs is a token-level three-class classification task. We used the same dataset as SpliceAI ^8^ to predict the splice sites of human pre-mRNAs, which consisted of 15,036 pre-mRNAs annotated by GENCODE V24lift37. The pre-mRNAs from chromosome 2, 4, 6, 8, 10-22, X, Y were used to fine-tune the model (13,384 transcripts,

130,796 pairs of splice sites), and the pre-mRNAs from chromosome 1, 3, 5, 7, 9 which did not have any paralogs were used for validation (1,652 transcripts, 14,289 pairs of splice sites). We used “bedtools” ^58^ to extract the pre-mRNA sequences between transcription start and end sites from the hg19/GRCh37 assembly.

We adopted the similar training strategy as SpliceAI ^8^ to fine-tune the model. The labels are predicted for a 768nt window each time, based on the sequence within the window and its flanking regions (128 nt on each side). In other words, the sequence of 1,024 nt was input into the model to classify the nucleotides in middle region (768 nt). Specifically, we first padded each pre-mRNA sequence with ‘N’ until the length became a multiple of 768, and further respectively padded 128 ‘N’s at the beginning and end of the sequence. The padded sequences were then split into blocks of 1024 nt, where the *i*_*j*ℎ_ block ranged from 768(*i* − 1) − 128 + 1 to 768*i* + 128, and the labels of nucleotides from 768(*i* − 1) + 1 to 768*i* were predicted during fine-tuning.

The cross-entropy loss function of predicting splice sites is denoted as formula (4):

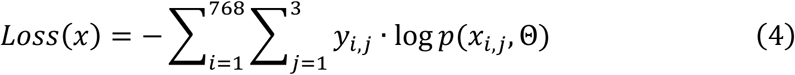

where *p*(*x*_*i*,_, Θ) represents the probability of predicting *i*_*j*ℎ_ nucleotide as class *j*, and *y*_*i*,_ represents whether *i*_*j*ℎ_ nucleotide belongs to class 푗. We selected the optimum combination of hyperparameters by grid searching (Supplementary table 1). As the training dataset is not balanced because most of the nucleotides are not splice sites, we use top-k accuracy and area under the precision-recall curve (PRAUC) as evaluation metrics ^8^. The top-k accuracy for the specified class is defined as follows: Suppose *k* positions belong to the class *j* in the testing set, the threshold of classification will be selected so that exactly *k* samples are predicted as class *j*. The top-k accuracy is computed as the proportion of *k* samples which are correctly classified. The top-k accuracy and PRAUC for predicting 5¢ and 3¢ splice sites are averaged for final evaluation.

#### 6.2. Prediction of translation efficiency

The 5¢ UTR sequences of mRNAs are input into LAMAR to predict the translation efficiencies of mRNAs. Prediction of the translation efficiency is a sequence-level regression task, only the embedding of 〈*cls*〉 token is fed into the task-specific head to fit the translation efficiency. We collected the translation efficiency of each mRNA from the Cao et al. dataset ^25^. The dataset contained human transcripts of which the expression and ribosome profiling were measured by RNA-seq and Ribo-seq in HEK293 cell line. We included the protein coding transcripts satisfying the following criterion: *RPKM*_*RNA*–*seq*_ ≥ 10 and *RPKM_Ribo–seq_* ≥ 10. Consistent with the methods of the original article ^25^, we extracted the 5¢ UTR sequences annotated by GENCODE V17 from hg19/GRCh37 assembly, and computed the translation efficiency of mRNAs by 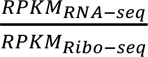. In all, we included 10,903 5¢ UTRs ranging from 5 nt to 1,024 nt in length. For 5¢ UTRs exceeding 1,024 nt, the segment of 1,024 nt at the 5¢ end of the start codon were extracted. We set the padding side as left in this fine-tuning task for better performance.

We select the mean squared error (MSE) loss to fine-tune the model to fit the translation efficiency, the function of which is denoted as formula (5):

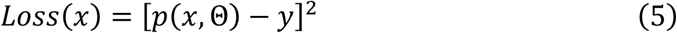

where *p*(*x*, Θ) represents the predicted translation efficiency of mRNA *x*, and *y* represents the observed value. The MSE and Spearman correlation coefficient between predicted translation efficiencies and observed ones are selected as metrics to evaluate the model.

#### 6.3. Prediction of degradation rates

We fine-tuned LAMAR with the 3¢ UTR sequences to predict the half-lives of mRNAs. Prediction of the half-lives is also a sequence-level regression task, only the embedding of 〈*cls*〉 is fed into the task-specific head to fit the half-lives detected from the experiment. We obtained the half-lives from the Zhao et al. dataset ^30^. The dataset measured the half-lives of 2,790 mRNAs with different 3¢ UTRs by the massive parallel reporter assays in BEAS-2B cell line and reported the half-lives of 1,967 3¢ UTR. We used these 3¢ UTR sequence and their half-lives to fine-tune the LAMAR. We selected the MSE loss (see formula 5) to fine-tune the model, and compute MSE and Spearman correlation coefficient for model evaluation.

To test whether the fine-tuned model accurately predicts the effects of 3¢ UTR variants on mRNA degradation, we selected the segment (67 nt) containing an AU-rich element in 3¢ UTR of CXCL2 which was saturated mutated as the positive control ^30^. The observed variant effect of each site was assessed by computing the difference between the steady-state mRNA levels of the wild-type and mutated sequences, and the predicted variant effect was measured by comparing the predicted half-lives of the wild-type and mutated sequences as the formula (6):

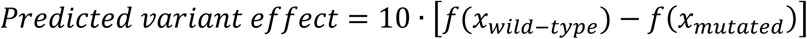

where *x*_*wild–type*_ means the native sequence, *x_mutated_* means the mutated sequences and *f*(*x*) means LAMAR-DR.

#### 6.4. Prediction of IRES

We fine-tuned LAMAR with different RNA classes to predict the IRES. Predicting the IRES is a sequence-level binary classification task, only the embedding of 〈*cls*〉 token is fed into the task-specific head to predict whether the RNA is IRES or not. We first collected the RNA sequences from IRESite ^39^, IRESbase ^40^ and RFAM ^41^ databases to build the training set. In total, the 1,901 experimentally validated IRES sequences and 2,026,606 non-IRES sequences were collected. To avoid the data leakage caused by sequence similarity, we used MMseqs2 ^59^ to cluster positive sequence and negative sequence respectively (identity=0.9, coverage=0.9), and obtained 903 positive clusters and 810,096 negative clusters. We sampled the representative sequences from the negative clusters so that the number of positive and negative sequences is the same and the distribution of sequence length is similar. We randomly selected 50% of the positive and negative clusters as the training set and the rest as the evaluation set. We repeated the experiment 10 times to measure the performance of the model.

We used the binary cross-entropy loss to fine-tune the model to classify the IRES. The loss function was denoted as:

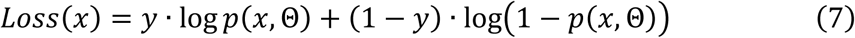

where *p*(*x*, Θ) represents the probability of predicting the RNA *x* as IRES, and *y* represents whether the RNA is IRES. We selected the optimum combination of hyperparameters by grid searching (Supplementary table 1). The area under the receiver operating characteristic (AUROC) was selected as evaluation metrics.

To validate the generalization of the fine-tuned model, we compared the predicted probabilities of IRESs and non-IRESs screened by three high-throughput experiments. For Weingarten et al. ^37^ dataset, the oligos derived from viral and human transcripts were inserted into bicistronic reporters and screened by fluorescence activated cell sorting (FACS). We filtered the oligos with promoter activity more than 0.2 and splicing activity less than -2.5, and labeled the sequences with IRES activities exceeding 600 as IRESs. Since a small part of oligos derived from IRESite database of which the sequences were used for training, we excluded them from the dataset to avoid data leakage. At last, 1,915 IRES and 17,047 non IRESs were included. The oligo library of Chen et al. ^36^ was the same with that of Weingarten et al., but was screened by circRNA reporter assays. Consistent with the original journal, we labeled the oligos with IRES activities exceeding 3.466 as IRESs. The oligos derived from IRESite database were filtered. At last, 9,185 IRESs and 11,353 non IRESs were included. For Yang et al. ^38^ dataset, the oligos were derived from whole human transcripts, inserted into circRNA reporters and screened by FACS. We labeled the oligos from post-sorting library as IRES and the difference set between the pre-sorting and post-sorting libraries as non-IRES. The oligos with lengths shorter than 50 nt or derived from IRESite database were filtered. To reduce the redundancy of sequenced oligos, we used MMseqs2 (identity=1.0, coverage=0.5, coverage mode=1, cluster mode=2) to cluster the oligos and selected the representative sequences. At last, 20,367 IRESs and 134,694 non IRESs were included.

### 7. Comparing the performance of LAMAR with baseline methods

We compared the performance of LAMAR with baseline methods including RNA-FM ^16^, spliceAI ^8^ and UTR-LM ^26^ across downstream tasks. The RNA-FM and UTR-LM were language models with architectures similar to LAMAR, pretrained on non-coding RNAs and 5¢ UTRs, respectively. For fine-tuning, we used the same prediction heads as LAMAR and initialized RNA-FM and UTR-LM with their pretrained weights. All model parameters were fine-tuned, with identical input/output data formats, training strategies and evaluation metrics as LAMAR. RNA-FM was fine-tuned for all downstream tasks, while UTR-LM was fine-tuned specifically for prediction of translation efficiency on 5¢ UTRs.

SpliceAI was a convolutional neural network designed to predict splicing from pre-mRNA sequences. It includes four architectures (SpliceAI-80, SpliceAI-400, SpliceAI-2k and SpliceAI-10k) which predicts splice sites within a sliding window of 5,000 nt using flanking sequences of different length (40nt, 200nt, 2,000 nt and 5,000 nt) on each side, respectively. To align with training length of LAMAR, we selected SpliceAI-400 and SpliceAI-2k as baseline methods for prediction of splice site. We directly applied their trained weights to predict splice sites from full-length pre-mRNAs, using the same evaluation metrics as LAMAR.

### 8. Identification of *cis*-elements regulating mRNA degradation

We used *in silico* mutation to identify *cis*-elements regulating RNA degradation. We mutated each nucleotide of 3¢ UTR segments and predicted the difference of half-lives between native and mutated sequences, which we named the perturbation score for each site. The perturbation score of the *i*^*j*ℎ^ nucleotide was defined by the formula:

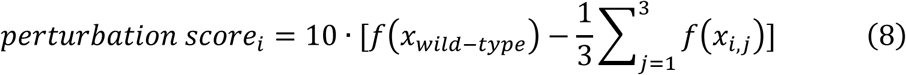

where, *x*_*wideȓtype*_ means the native sequence, *x*_*i*,_ means the sequence whose *i*^*th*^ site is mutated to *j^jh^* class of nucleotide and *f*(*x*) means LAMAR-DR. To increase the stability of prediction, we used ten sub-models to predict half-lives of native and mutated sequences and averaged the perturbation scores as the final result.

The perturbation scores could represent the regulatory effect of the elements. The negative perturbation score implied that the mutation breaks the destabilizing element and extend the half-life of mRNA. The positive perturbation scores implied that the mutation breaks the stabilizing element and shorten the half-life of mRNA.

To validate the rationality of perturbation score, we selected 150 reported regulatory elements as positive controls ^30^. These elements were screened based on Tet-Off system, and their regulatory effects were measured by the difference in the fraction of mRNA remaining after 4 h of doxycycline treatment between mutant and wild-type clones. We averaged the perturbance score of each site in the element to predict its regulatory effect for further studies.

As for identification of *cis*-elements with strong regulatory effects from the training set, we averaged the perturbation scores of six consecutive nucleotides to represent the regulatory effects of the hexamers. The hexamers with highest and lowest 1% perturbation scores and frequency more than 10 were selected. The hexamers were further multiple aligned by Cluster Omega ^62^ and clustered by pairwise distance to a UPGMA phylogenetic tree. We plotted the motifs for each cluster by R package ggseqlogo ^63^.

### 9. *In silico* screening of IRES

We used the fine-tuned model LAMAR-IRES to predict the putative IRESs and validated their activities by experiments. We first downloaded the genome sequences and GenBank annotations of positive-sense single-stranded RNA viruses (SARS-CoV-2 excluded) from NCBI virus database. The annotated 5¢ UTRs were thought as putative IRESs and extracted from the genomes. If there was no annotated 5¢ UTR, the segment before the first CDS would be included. Further, we excluded the 5¢ UTRs containing too many ‘N’s and removed the duplicates. We used prokka ^64^ to predict the putative coding regions in 5¢ UTRs and deleted these sequences from the candidates. The remaining 5¢ UTRs with the length ranged from 500∼1500 nt were clustered twice and 2,067 representative sequences of clusters were included. We preferentially selected 305 5¢ UTRs of the viruses which infect vertebrates for *in silico* screening.

We used 10 sub-models of LAMAR-IRES to score the tested 5¢ UTRs and categorized them into 3 groups by the averaged probabilities: non-IRESs (probability<0.5), low-activity IRESs (0.5≤probability<0.9) and high-activity IRESs (0.9≤probability<1). To validate the correlation between predicted probability and IRES activity, the efficiency of 5¢ UTR promoting circRNA translation was measured. Specifically, the sequences were inserted into the *in-vitro* synthesized Fluc-coding circRNA and transfected into 13 kinds of cell lines (A204, A549, AGS, C2C12, HCT116, HepG2, HFF1, JURKAT, Neruo-2a, RAW264.7, SY5Y, THP-1 and U87) with 96-well plate. The mock and modified Fluc-coding mRNA (CATUG Bio) were transfected into cells as controls in each batch. Each sample had two replicates. At 24 h after transfection, cells were collected and lysed in the passive lysis buffer. The protein expression level was measured via luminescence intensity. The mock transfection was used as an internal control and its protein expression was subtracted.

The relative protein expression of circRNA and modified mRNA was computed and the final IRES activity was defined by the following formula:

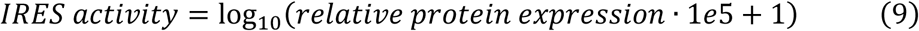

## Data availability

The datasets used for pretraining and fine-tuning are all derived from databases and previous published studies. Here we include the official websites. RefSeq (Release 221): https://ftp.ncbi.nlm.nih.gov/refseq/release/vertebrate_mammalian/; RNACentral (Release 23): https://ftp.ebi.ac.uk/pub/databases/RNAcentral/releases/23.0/sequences/; NCBI Genome: https://ftp.ncbi.nlm.nih.gov/genomes/refseq/vertebrate_mammalian/assembly_summary.txt; NCBI Virus: https://ftp.ncbi.nlm.nih.gov/genomes/refseq/viral/assembly_summary.txt; SpliceAI dataset: https://basespace.illumina.com/s/5 u6ThOblecrh. RFAM (Release 14.9): https://rfam.org/search?q=IRES%20AND%20entry_type:%22Family%22. IRESite: http://www.iresite.org; IRESbase: http://reprod.njmu.edu.cn/cgi-bin/iresbase/index.php. The translation efficiency dataset was obtained from Cao J, et al. and the degradation dataset was obtained from Zhao W, et al. Three IRES datasets (Figure 6D) were obtained from Weingarten-Gabbay S et al, Chen C-K et al and Yang Y et al. Source data are provided with this paper.

## Code availability

The scripts used in this study are available in Github repository (https://github.com/rnasys/LAMAR).

## Acknowledgments

This work is supported by the National Key Research and Development Program of China (2021YFA1300503), the Strategic Priority Research Program of Chinese Academy of Sciences (XDB38040100) and the National Natural Science Foundation of China (32030064 and 32250013) to Z.W. This work is also supported by Shanghai Science and Technology Innovation Action Plan (23JS1401500) and Shanghai Municipal Science and Technology Major Project (GTP) to G.Z.

## Figures and figure legends

**Supplementary Figure 1.**
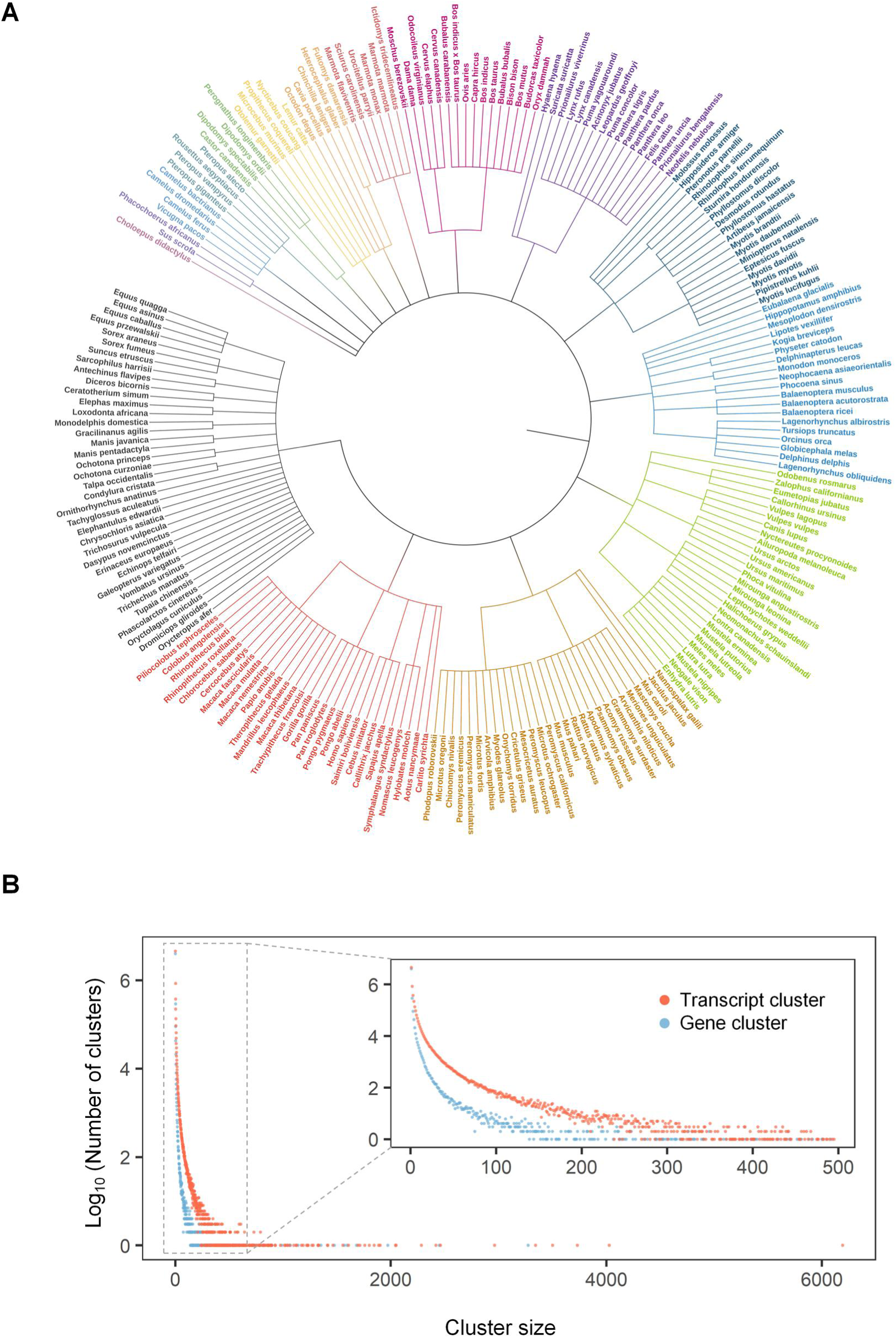
Summary of pretraining dataset. **(A)** Phylogenetic tree of mammals in pretraining dataset. The 225 mammals whose genes/transcripts were used in pretraining were clustered into 15 suborders by the phylogenetic tree. The mammals belong to different suborders are represented by different colors. **(B)** The number of gene/transcript cluster are plotted against the cluster size. Inset plot shows distribution of cluster size below 500.

**Supplementary Figure 2.**
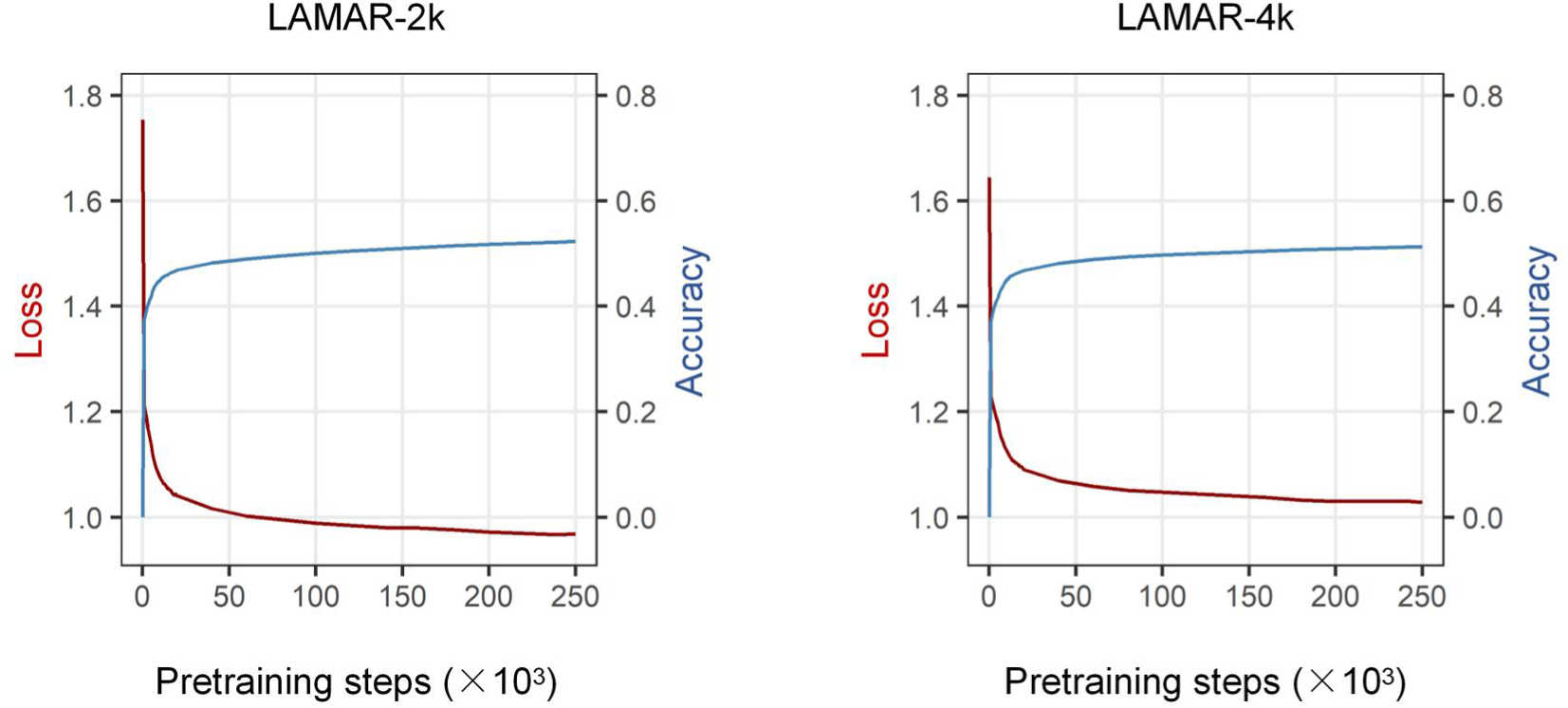
Evaluation of the pretraining steps. The loss curve for LAMAR-2k and LAMAR-4k models during the pretraining (rea lines). The accuracy of these models in predicting the masked tokens were also plotted in each step of pretraining (blue lines).

**Supplementary Figure 3.**
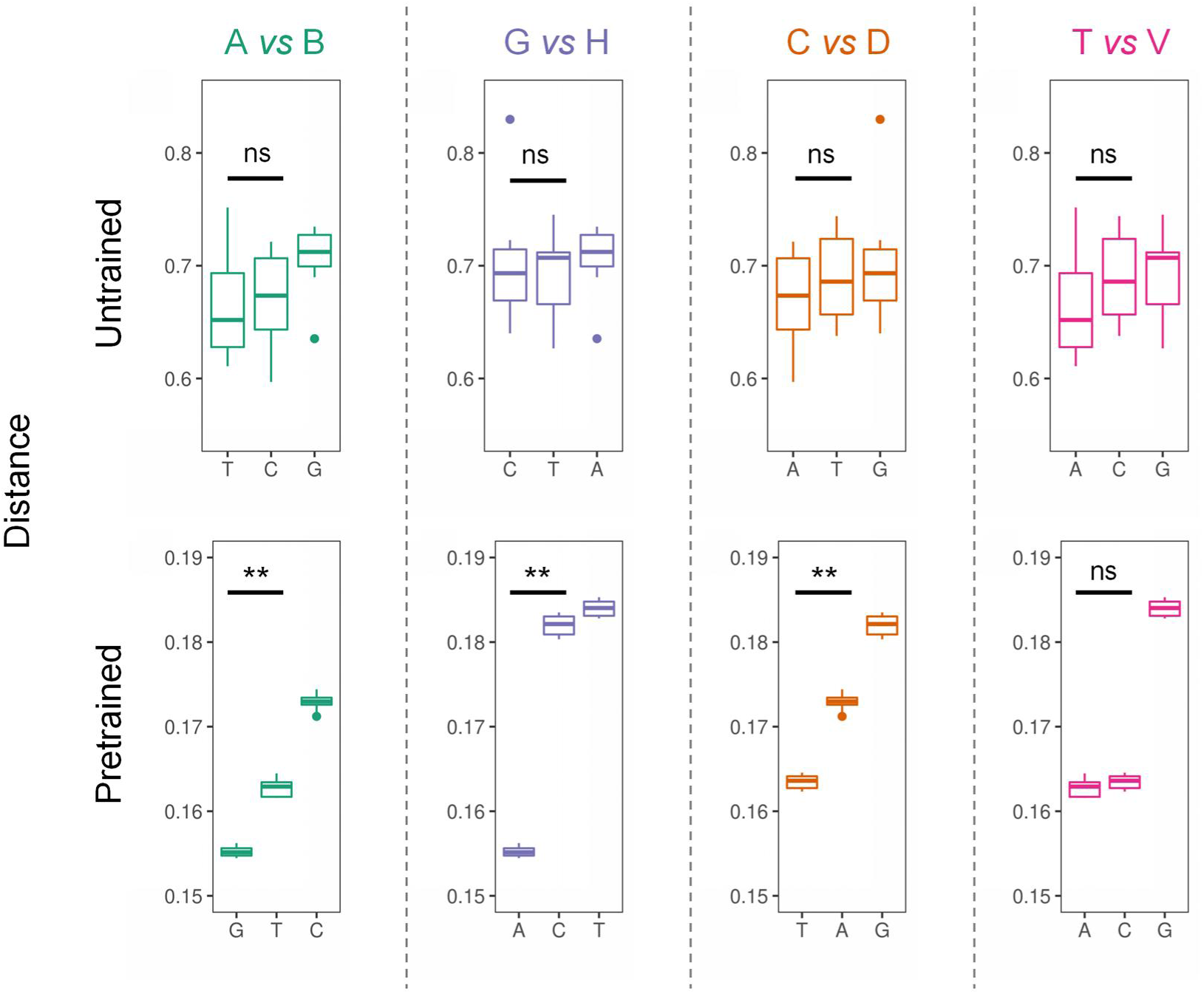
The distances between the nucleotide embeddings. The cosine distances between the nucleotide embeddings from untrained model (top) and pretrained model (bottom). *P* values were calculated with two-sided Wilcoxon rank sum test. **, *P* value < 0.01; ns, not significant.

**Supplementary Figure 4.**
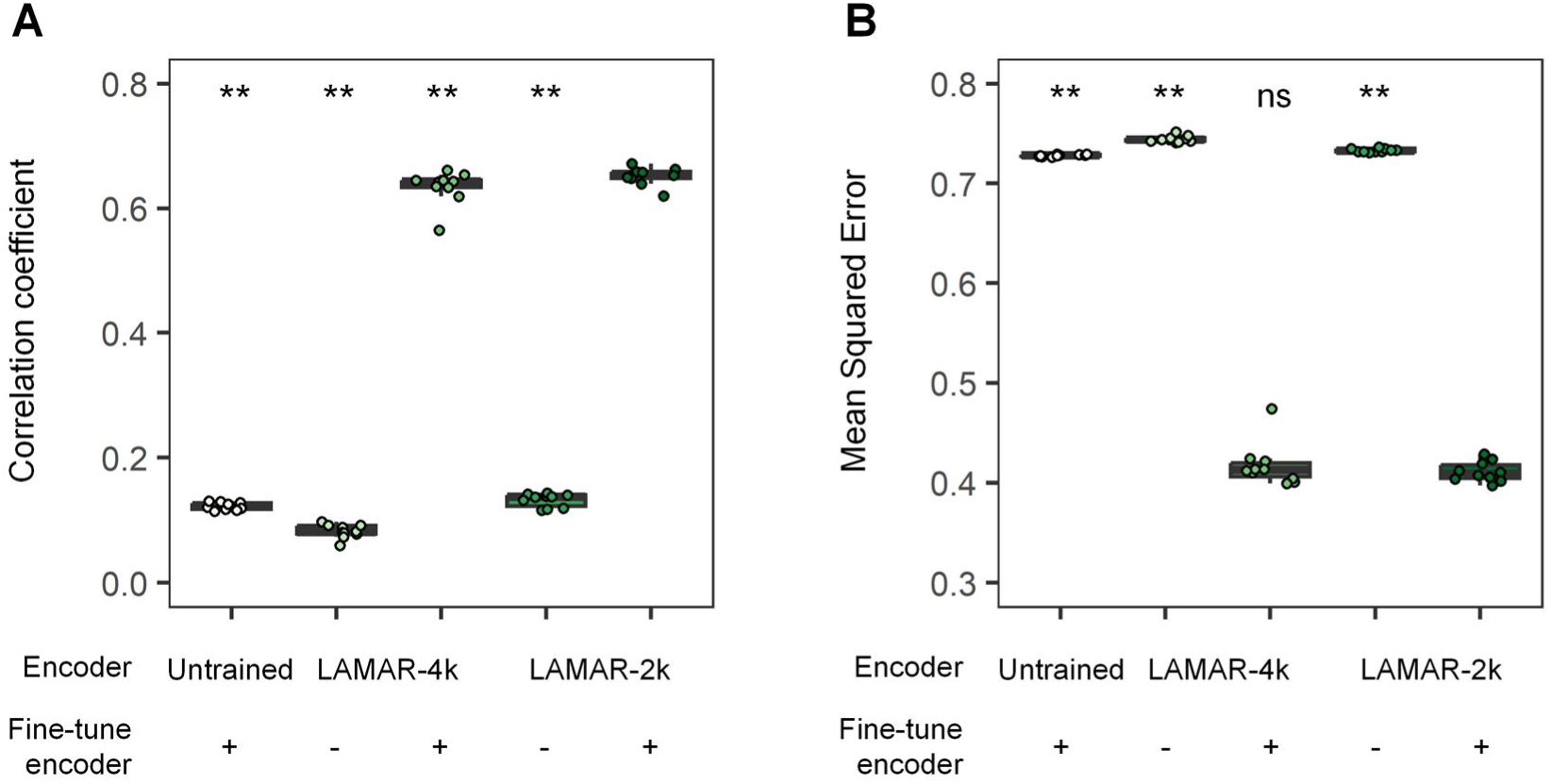
The performance of LAMAR fine-tuned with different strategies in predicting translation efficiency. The Spearman correlation coefficient **(A)** and mean squared error **(B)** between observed and predicted translation efficiency for LAMAR fine-tuned with different strategies. *P* values were calculated with Wilcoxon signed rank test. **, *P* value < 0.01; ns, not significant.

**Supplementary Figure 5.**
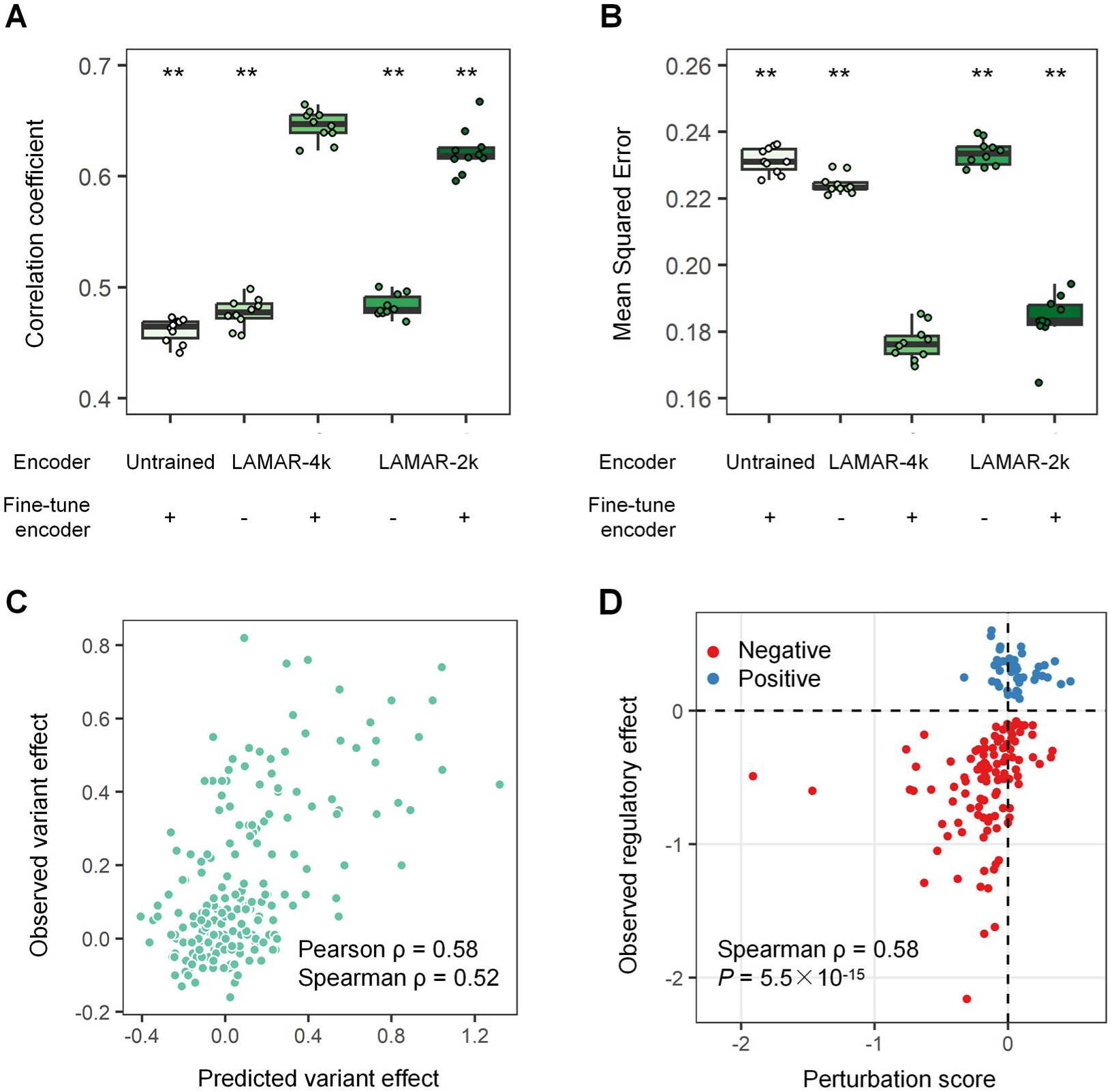
The performance of LAMAR fine-tuned with different strategies in predicting degradation rate. The Spearman correlation coefficient **(A)** and mean squared error **(B)** between observed and predicted half-life for LAMAR fine-tuned with different strategies. **(C)** The scatter plot depicting the predicted (x axis) and observed variant effect (y axis) of each position in 3′ UTR segment of CXCL2. **(D)** The scatter plot depicting the perturbation score (x axis) and observed regulatory effect (y axis) of the short elements which positively and negatively regulating the half-live of mRNA. *P* values were calculated with Wilcoxon signed rank test. **, *P* value < 0.01.

**Supplementary Figure 6.**
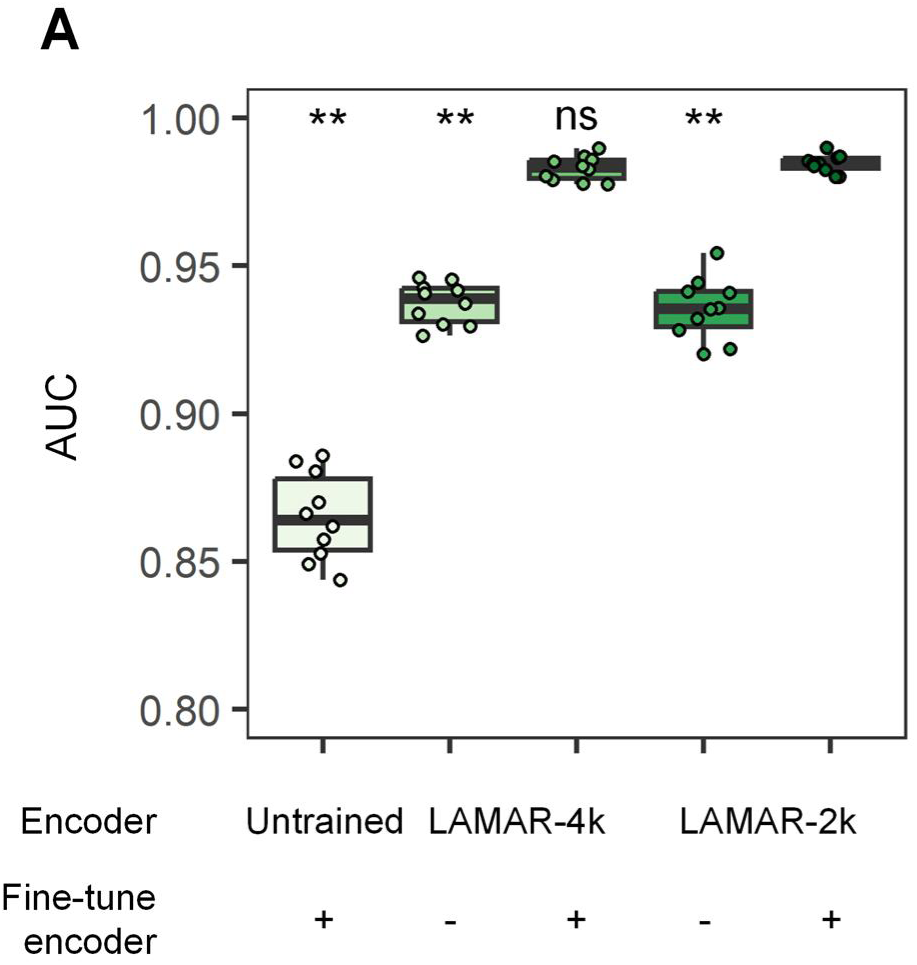
The performance of LAMAR fine-tuned with different strategies in predicting IRES. The AUC of LAMAR in classifying IRES, evaluated under different fine-tuning strategies. *P* values were calculated with Wilcoxon signed rank test. **, *P* value < 0.01; ns, not significant.

